# MDL-001: an Oral, Direct-Acting Broad-Spectrum Antiviral for Influenza-Like Illness (ILI) and Chronic Hepatitis In Vivo

**DOI:** 10.1101/2025.01.13.632836

**Authors:** Virgil Woods, Tyler Umansky, Sean M Russell, Asha Goodman, Michael Bobardt, Briana L McGovern, Romel Rosales, M Luis Rodriguez, Harm van Bakel, Emilia Mia Sordillo, Viviana Simon, Adolfo García-Sastre, Kris M White, Philippe Gallay, Davey Smith, Daniel Haders

## Abstract

**Background/Objectives:** Endemic viral respiratory illnesses caused by influenza, RSV, and coronaviruses impose a global disease burden of hundreds of millions of infections and hundreds of thousands of deaths annually. Chronic hepatitis B and C infections persist in 254 million and 58 million people worldwide, respectively. No approved therapy addresses these viruses simultaneously. MDL-001 is an oral, direct-acting, broad-spectrum antiviral targeting the Thumb-1 allosteric site of viral polymerases.

**Methods:** Here we evaluated the antiviral activity of MDL-001 in vitro in influenza A/B, RSV, SARS-CoV-2, endemic human coronaviruses, and hepatitis B/C/D. We next evaluated MDL-001 antiviral activity in vivo in H1N1, SARS-CoV-2, HCV, and HBV. MDL-001’s in vivo efficacy was compared head-to-head versus standards of care including oseltamivir, remdesivir, sofosbuvir, and tenofovir alafenamide, or through literature comparison with nirmatrelvir and molnupiravir. Lastly, we evaluated the pharmacokinetics, metabolism, tolerability, and safety profile of MDL-001.

**Results:** Here, we report MDL-001 inhibits influenza A/B, RSV, SARS-CoV-2, endemic human coronaviruses, and hepatitis B/C/D viruses with nanomolar EC_90_ potency in vitro. MDL-001 demonstrated equivalent in vivo efficacy to oseltamivir against mouse-lethal influenza A virus infection, reducing lung viral load by 2.6 log_10_ and preventing weight loss and mortality. MDL-001 demonstrated equivalent symptom reduction to subcutaneous remdesivir after SARS-CoV-2 infection. MDL-001 reduced SARS-CoV-2 lung viral titers by 2.9 log_10_, superior to literature reported 1.4 log_10_ and 1.0 log_10_ reductions for nirmatrelvir and molnupiravir, respectively. MDL-001 reduced HCV viremia by 3.3 log_10_, equivalent to sofosbuvir. Oral MDL-001 reduced plasma HBV surface antigen by 2.5 log_10_, superior to tenofovir alafenamide from Day 8 through Day 21. MDL-001 demonstrated an in vitro EC_50_ of 79.4 nM against infection with RSV strain A2, a 416-fold improvement over literature-reported ribavirin. Oral pharmacokinetics studies demonstrate MDL-001 is rapidly absorbed and partitioned into Lung (K_p_=39-52) and Liver (K_p_=71-104) tissues. MDL-001 produced no treatment-related tolerability adverse events across 376 animals and showed no hERG, Ames genotoxicity, or micronucleus safety issues.

**Conclusions:** These findings support MDL-001 as a broad-spectrum, direct-acting non-nucleoside antiviral clinical candidate.

## 1. Introduction

Each winter clinicians face a recurring tripledemic of influenza, RSV, and SARS-CoV-2. These viral respiratory illnesses (VRIs) impose a global disease burden of hundreds of millions of infections and hundreds of thousands of deaths annually [1]. In a non-pandemic year in the U.S., adults experience two to six VRIs and children approximately six to eight, with economic costs exceeding $100 billion [2]. There are currently no single-agent antivirals approved to treat this annual convergence of the tripledemic respiratory viruses.

Chronic viral hepatitis is a health crisis of enormous proportions. An estimated 5–15 million people worldwide have both HBV and HCV infections [3–4]. Most will never be diagnosed. Only 13% of HBV infections and 36% of HCV infections are diagnosed globally. Fewer than 1% of co-infected patients know their status [5]. Current treatments for chronic hepatitis C possess a critical and potentially fatal vulnerability. Every FDA-approved HCV direct-acting antiviral (DAA) carries a black box warning for the risk of hepatitis B virus (HBV) reactivation in patients with current or prior HBV infection [6]. This FDA-mandated warning applies to DAA regimens such as Epclusa, Harvoni, and Mavyret. HBV reactivation in HCV co-infected patients can lead to fulminant hepatitis, liver failure, or death. The ideal therapeutic for HCV/HBV co-infection would be an oral, well-tolerated, direct-acting antiviral with activity against both viruses through a single mechanism of action. This agent would eliminate the need for fragmented drug regimens, reduce the risk of HBV reactivation during HCV clearance, and simplify treatment delivery. No approved single-agent therapy directly addresses both HCV and HBV. No major clinical trial in the HBV functional cure pipeline specifically enrolls patients coinfected with HCV. This population remains systematically understudied and underserved.

Together, these gaps point to a persistent and largely unaddressed need for ILI, chronic hepatitis and pandemic preparedness: a single agent with broad-spectrum, direct-acting antiviral activity against RNA viruses and viruses with an RNA step. No oral non-nucleoside direct-acting antiviral has demonstrated activity across multiple RNA virus families. The United States’ National Institute of Allergy and Infectious Diseases (NIAID) Target Product Profiles (TPP) for Antivirals defines this gap explicitly for pandemic preparedness [7–8]. The ideal countermeasure is orally bioavailable, active across viral families, suitable for outpatient use, and stable enough for stockpiling before an outbreak. No approved antiviral meets these criteria.

Conventional scientific wisdom holds that allosteric sites on viral polymerases diverge too widely to enable the development of broad-spectrum direct-acting antivirals (DAAs). We overturned this assumption by demonstrating conservation of the RNA-dependent RNA polymerase (RdRp) Thumb-1 allosteric pocket across viral families [9]. The Thumb-1 site governs an essential mechanism in viral replication. The Thumb-1 pocket interacts with the viral Λ1-loop to control an indispensable conformational change required for polymerase initiation. Engagement at this site traps the polymerase in a catalytically incompetent state without competing with the active site for substrate. Viruses cannot abandon this mechanism without sacrificing their replicative fitness. Beclabuvir is the only approved Thumb-1 inhibitor [10]. Beclabuvir was developed for hepatitis C, and preclinical selectivity studies demonstrated that it was inactive (EC_50_ >14 µM) against a diverse panel of non-HCV viruses, including poliovirus, rhinovirus, coronavirus, coxsackievirus, influenza, and HIV [11]. A single small molecule that can engage Thumb-1 with the potency, oral pharmacokinetics, and tolerability needed for in vivo efficacy across multiple viral families has not been demonstrated. The discovery of the broad-spectrum Thumb-1 pocket, the discovery of MDL-001 and resistance, biochemical and cellular confirmation of MDL-001 engagement of the Thumb-1 pocket and antiviral activity has been reported previously.

Here, we report the preclinical PoC characterization of MDL-001 (pipendoxifene), an oral non-nucleoside small molecule with nanomolar antiviral activity across multiple virus families. The ChemPrint geometric deep-learning platform identified MDL-001 within the GALILEO drug discovery pipeline [12–14]. The GALILEO platform and ChemPrint model used to discover MDL-001 have been peer-reviewed and published previously [15]. We demonstrate nanomolar in vitro activity against influenza A and B, RSV, SARS-CoV-2 and variants of concern, endemic human coronaviruses, and hepatitis B, C, and D viruses. Oral MDL-001 achieves in vivo efficacy equivalent to oseltamivir for influenza A symptom protection and lung viral reduction, subcutaneous remdesivir for SARS-CoV-2 symptom protection, and sofosbuvir for HCV viremia. Oral MDL-001 exceeds published SARS-CoV-2 lung-titer benchmarks for nirmatrelvir and molnupiravir [16–17]. In vitro, MDL-001 exceeds the published RSV A2 potency benchmark for ribavirin [18]. Oral MDL-001 also exceeds tenofovir alafenamide for reduction of plasma HBV surface antigen. Oral pharmacokinetics data demonstrate MDL-001 is rapidly absorbed and distributed, specifically partitioning into the Liver and Lung. Tolerability evaluation across 376 animals reveals no treatment-related adverse findings. No hERG, Ames genotoxicity, or micronucleus safety issues were observed. These data establish MDL-001 as a clinical candidate that meets the NIAID Target Product Profile for a broad-spectrum direct-acting antiviral. MDL-001 clinical development has the potential to define a path to outpatient deployment against novel viral respiratory pandemic threats, the annual endemic respiratory tripledemic and the global chronic hepatic viral disease crisis.

## 2. Results

### In vitro Cross-Family Antiviral Activity of MDL-001

MDL-001 was evaluated in vitro against a panel of respiratory and hepatic viruses that utilize right handed polymerases or reverse transcriptases to evaluate antiviral activity. MDL-001 inhibited influenza A and B, respiratory syncytial virus, SARS-CoV-2 and variants of concern, endemic human coronaviruses, and hepatitis B, C, and D viruses with nanomolar potency. EC_50_ values ranged from 27 to 263 nM across all viruses tested. EC_90_ values ranged from 47 to 595 nM across the respiratory panel and from 88 to 460 nM across the hepatic panel ( **Fig. 1; Supplementary Table S1**).

**Figure 1.**
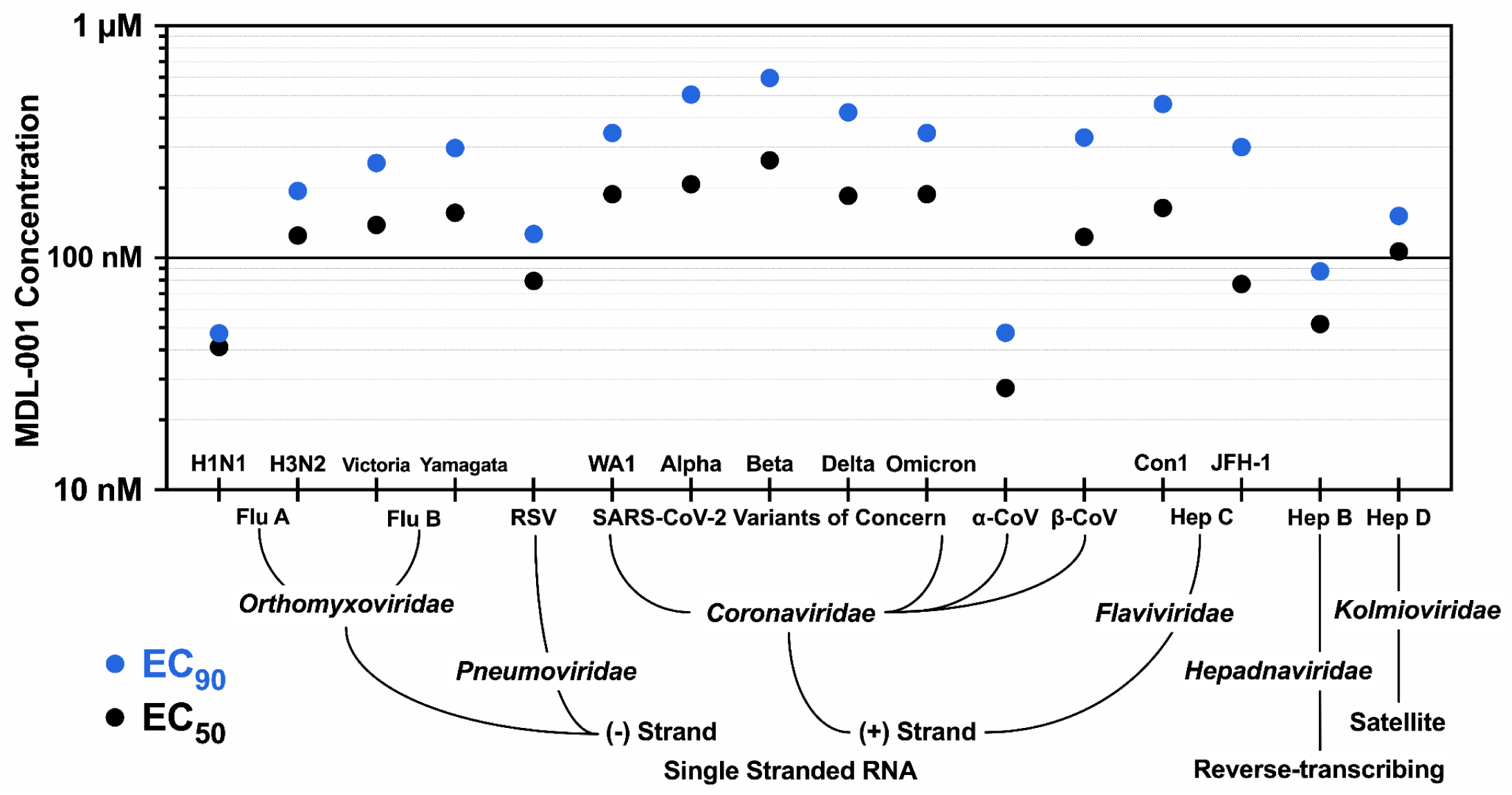
Broad-spectrum antiviral activity of MDL-001 across hepatic and respiratory viruses. Potencies (EC_50_, black; EC_90_, blue) for MDL-001 are plotted on a logarithmic y-axis. Values were obtained from concentration-response curves in the indicated cell-based infection assays. Assay parameters are listed in **Supplementary Table S1**.

### In vivo Cross-Family Efficacy of Oral MDL-001

#### Oral MDL-001 Establishes Equivalency to Oseltamivir in an Influenza A Mouse Challenge

MDL-001 and the clinical equivalent dose of oseltamivir [19] were evaluated against mouse-lethal H1N1 influenza A virus challenge in vivo. The two experimental arms and the vehicle arm were compared to evaluate the superiority, equivalence, or inferiority of MDL-001 to oseltamivir in reducing symptomatic disease progression and lung viral titer at Day 5. Symptomatic disease progression was measured as body-weight loss and mortality. Vehicle-treated mice lost body weight progressively over the observation period ( **Fig. 2A**). Mean body weight declined to approximately 91% of baseline by Day 5 post-infection. All three vehicle-treated mice died in the PM of Day 5 p.i. All MDL-001-treated and oseltamivir-treated mice survived to Day 5 p.i. and gained weight from Day 1 onward, reaching approximately 103% of baseline by Day 5 AM. Oral MDL-001 was equivalent to oseltamivir for body-weight preservation (*P = 0.81; 90%CI −1.0 to +1.9%*) on Day 5 p.i. Oral MDL-001 reduced lung viral titers by 2.6 log_10_, equivalent to oseltamivir on Day 5 p.i. *(P = 0.14; 90%CI −0.3 to 0.0 log_10_;* **Supplementary Table S2**).

**Figure 2.**
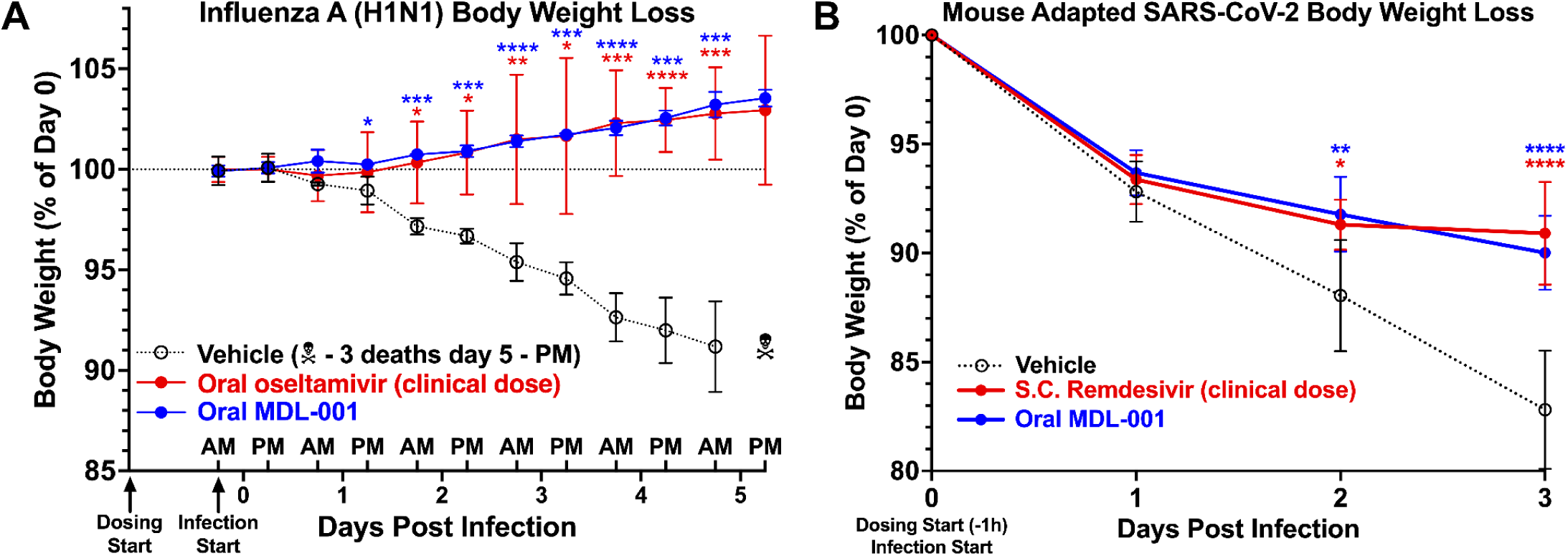
In vivo efficacy of oral MDL-001 against respiratory viruses. **(A)** Balb/c mice (n = 3 per group) were treated orally with MDL-001, the clinically equivalent oral dose of oseltamivir [19], or vehicle 24 hours prior to being intranasally infected with 1.0 × 10^5^ PFU of Influenza A/H1N1/PR8-luc. Twice-daily body weight measurements are expressed as percentage of Day-0 baseline from Day 0 through Day 5 p.i.; vehicle group n = 0 at Day 5 PM due to mortality. At Day 5, the between-arm difference in body weight was within the pre-specified ±5% equivalence margin, with a 90% confidence interval of −1.0 to +1.9% of baseline (equivalent to oseltamivir, P = 0.81).**(B)** 129/S mice (n = 9 per group) were treated orally with MDL-001, subcutaneously with the clinically equivalent dose of remdesivir [20], or vehicle 1 hour prior to intranasal infection with 2.5 × 10^4^ PFU of MA-SARS-CoV-2. Dosing continued for 3 days. Animal weights were monitored daily. At Day 3, the between-arm difference in body weight was within the pre-specified ±5% equivalence margin, with a 90% confidence interval of −3.1 to +1.3% of baseline (equivalent to remdesivir, P = 0.47). **(A-B)** Points denote group means; error bars denote 95% CI. Significance of difference from vehicle control was tested by two-way ANOVA (**\***P < 0.05, **\*\***P < 0.01, **\*\*\***P < 0.001, and **\*\*\*\***P < 0.0001).

#### Oral MDL-001 Establishes Equivalency to Remdesivir in a SARS-CoV-2 Mouse Challenge

MDL-001 and the clinical equivalent dose of subcutaneous remdesivir [20] were evaluated for symptomatic attenuation of SARS-CoV-2 infection in vivo. The experimental arms and the vehicle arm were compared to evaluate the superiority, equivalence, or inferiority of MDL-001 to remdesivir in reducing body-weight loss at Day 3. Mean body weight declined to approximately 82% of baseline by Day 3 p.i. in the vehicle group. Disease-associated weight loss was significantly attenuated by MDL-001 at Days 2 p.i. and 3 p.i. Oral MDL-001 was equivalent to remdesivir for body-weight preservation on Day 3 (*P = 0.47; 90%CI −3.1 to +1.3%;* **Fig. 2B**).

MDL-001 was evaluated for reduction of SARS-CoV-2 viral titers in vivo. Oral MDL-001 reduced lung viral titers at Day 3 p.i. by 2.9 log_10_ versus vehicle (**Supplementary Table S3**). MDL-001 and the literature-reported clinical equivalent doses of nirmatrelvir and molnupiravir were compared to evaluate the superiority, equivalence, or inferiority of MDL-001 to these literature reported values. The 2.9 log_10_ reduction reported here is superior to the 1.4 log_10_ reduction reported in the literature for the clinically equivalent dose of nirmatrelvir (*P* = 0.023) [16] and the 1.0 log_10_ reduction reported in the literature for the clinically equivalent dose of molnupiravir (*P* = 0.008) [17].

#### Oral MDL-001 Establishes Equivalency to Sofosbuvir in an HCV Mouse Challenge

MDL-001 and the clinical equivalent dose of oral sofosbuvir [21] were evaluated against HCV infection in vivo. The experimental arms and the vehicle arm were compared to evaluate the superiority, equivalence, or inferiority of MDL-001 to sofosbuvir in reducing HCV RNA over 28 days. Oral MDL-001 reduced plasma HCV RNA by 3.3 log_10_ at Day 28. Oral MDL-001 was equivalent to sofosbuvir at Day 28 (*P = 0.75; 90%CI −0.2 to +0.1 log_10_;* **Fig. 3A-C**). Oral MDL-001 reached equivalency with sofosbuvir by Day 21 and remained equivalent through Day 28 (**Supplementary Table S4**). Both arms approached the assay lower limit of quantification at Day 28.

**Figure 3.**
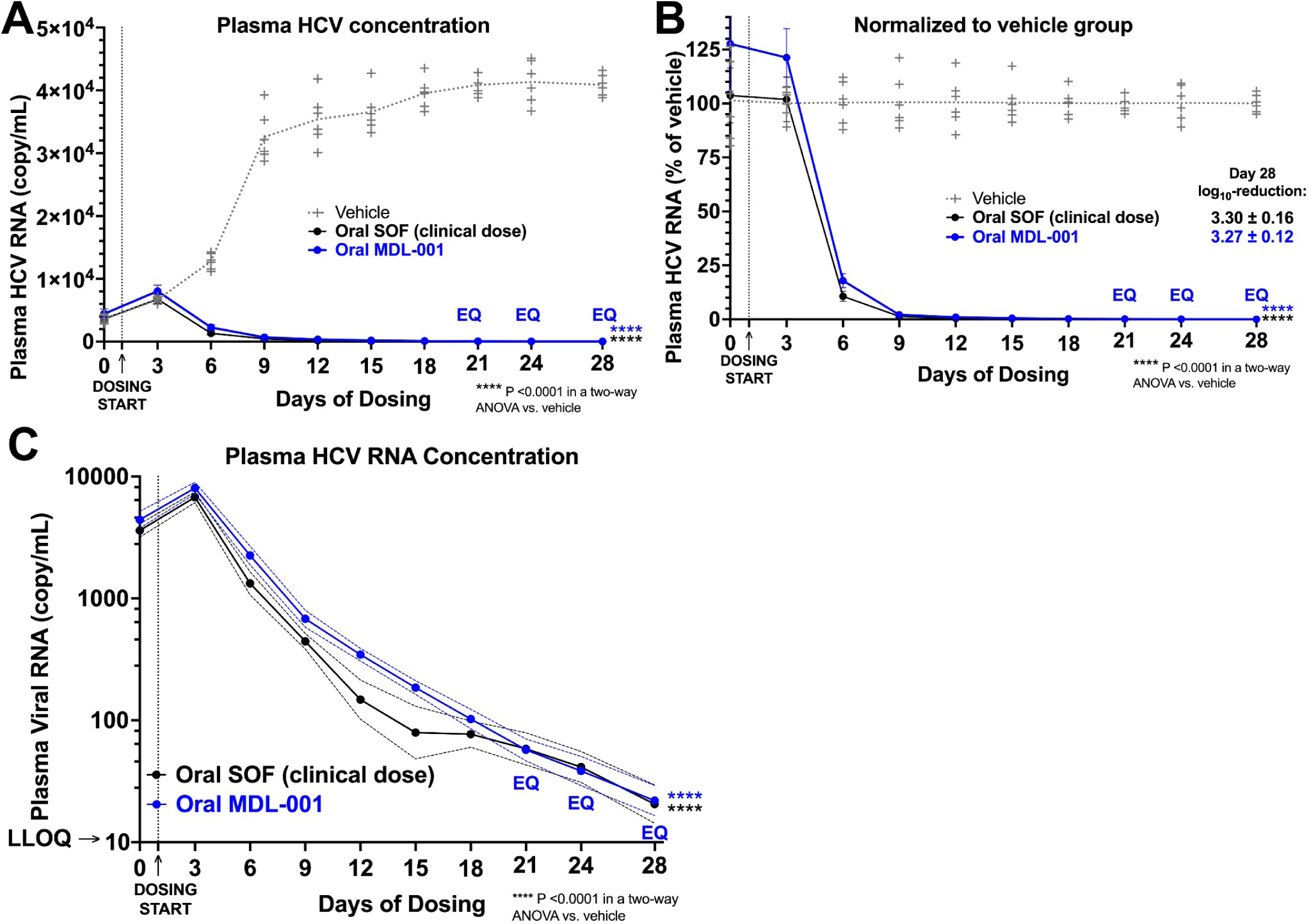
In vivo efficacy of oral MDL-001 against hepatitis C virus. **(A)** Humanized-liver mice (n = 6 per group) were infected intravenously on Day −2 with 1 × 10^6^ IU HCV JFH1/J6 and treated orally with MDL-001 or the clinically equivalent dose of sofosbuvir (SOF) [21], or vehicle starting on Day 1. Plasma viral genome copies per mL (qPCR) were measured one day before dosing and every three days thereafter until Day 28. **(B)** The same plasma data expressed as a percent of the contemporaneous vehicle mean. **(C)** The same plasma data on a logarithmic axis with vehicle omitted for visual clarity. Dashed lines denote the bounds of the 95% CI. **(A-B)** Points denote group geometric means; error bars denote 95% CI. Significance of reduction from vehicle control was tested by two-way ANOVA (****P < 0.0001). EQ denotes days where MDL-001 reached statistical equivalence with sofosbuvir. (**Supplementary Table S4**).

#### Oral MDL-001 Establishes Superiority to Tenofovir Alafenamide in an HBV Mouse Challenge

MDL-001 and oral tenofovir alafenamide (TAF) were evaluated against HBV infection in vivo. The two experimental arms and the vehicle arm were compared to evaluate the superiority, equivalence, or inferiority of MDL-001 to tenofovir alafenamide in reducing plasma HBsAg over 28 days. Oral MDL-001 produced a greater reduction in plasma HBsAg than TAF at every sampling timepoint from Day 8 through Day 21 (Day 8, P = 0.004; Day 19, P < 0.0001; Day 21, P = 0.006; **Fig. 4B-C; Supplementary Table S5**). The separation was largest at Day 19, where MDL-001 reduced plasma HBsAg by 1.5 log_10_ against 1.20 log_10_ for TAF. Oral MDL-001 reduced plasma HBsAg by 2.5 log_10_ at Day 28, at which point both arms had converged on the assay lower limit of quantification (P = 0.36; **Fig. 4A-C**). Both arms reduced plasma HBsAg significantly below the vehicle from Day 2 onward. Plasma human albumin levels were preserved across all arms.

**Figure 4.**
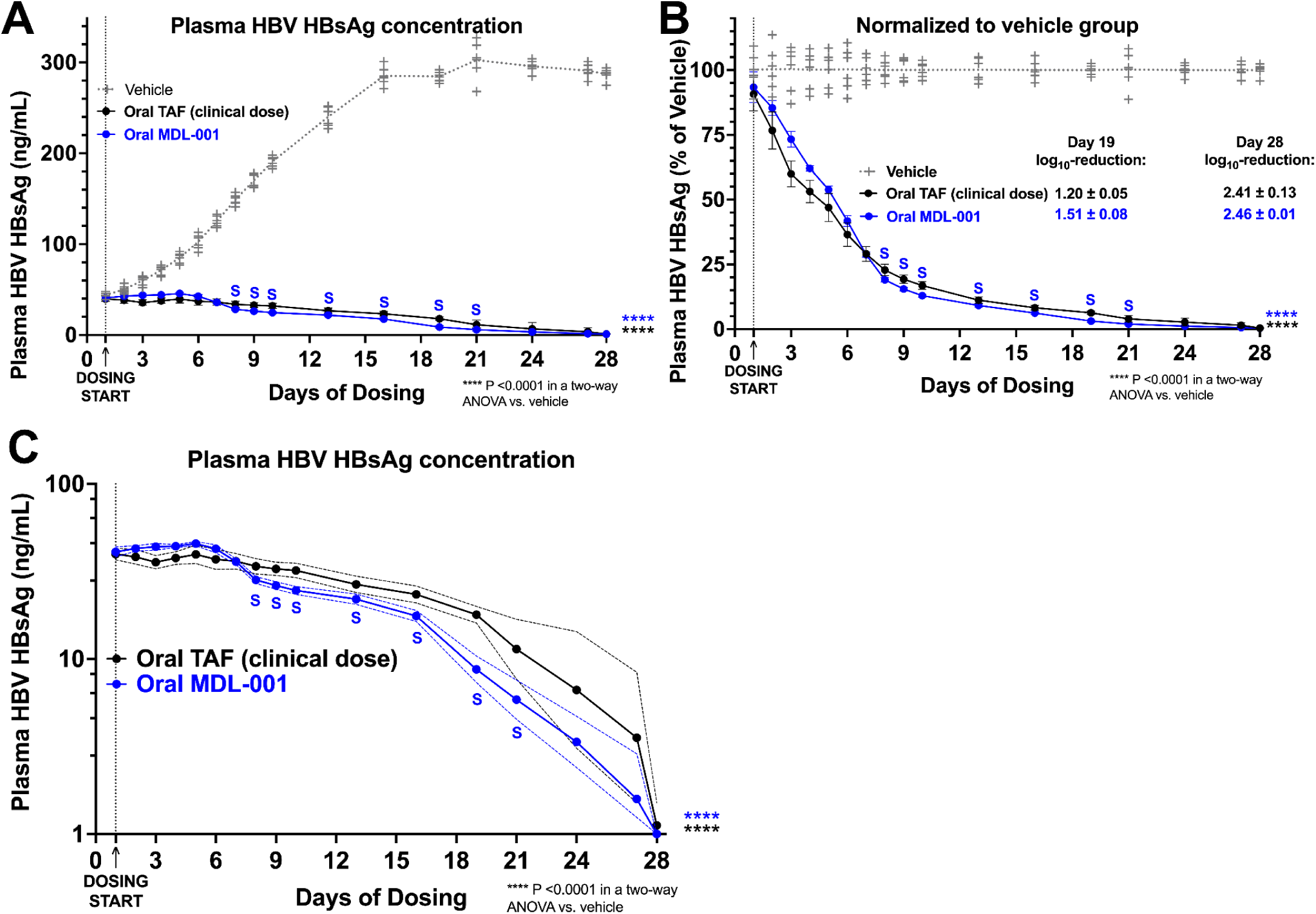
In vivo efficacy of oral MDL-001 and tenofovir alafenamide against hepatitis B virus. **(A)** Humanized-liver mice (n = 6 per group) were infected intravenously on Day −2 with 1 × 10^7^ genome equivalents of HBV (AD38 strain) and treated orally with MDL-001, TAF, or vehicle starting on Day 1. Plasma HBsAg concentration (ng/mL) was measured at seventeen timepoints through Day 28. **(B)** The same data expressed as a percent of the contemporaneous vehicle mean. **(C)** The same data on a logarithmic axis with vehicle omitted for visual clarity. Dashed lines denote the bounds of the 95% CI. **(A-C)** Points denote group geometric means; error bars and dashed lines denote 95% CI. Significance of reduction from vehicle control was tested by two-way ANOVA (****P < 0.0001). Blue S symbols denote days where MDL-001 reached statistical superiority over TAF. (**Supplementary Table S5**).

### Pharmacokinetics, Metabolism, Tolerability and Safety Profile of MDL-001

#### Pharmacokinetics: Oral MDL-001 Rapidly Absorbed and Partitioned Extensively into Target Tissues

A single-dose oral pharmacokinetic study characterized MDL-001 exposure in plasma, lung, and liver at three dose levels (**Fig. 5A-B; Supplementary Table S6**). Oral MDL-001 in suspension was rapidly absorbed and partitioned extensively into both target tissues. Tissue concentrations exceeded plasma at all sampled timepoints across all doses. Lung K_p_ ranged from 39 to 52 by C_max_ and from 38 to 76 by AUC0–24h across the 25 to 75 mg/kg dose range (**Fig. 5A**). Liver K_p_ ranged from 71 to 104 by C_max_ and from 65 to 91 by AUC0–24h across the same range (**Fig. 5B**).

**Figure 5.**
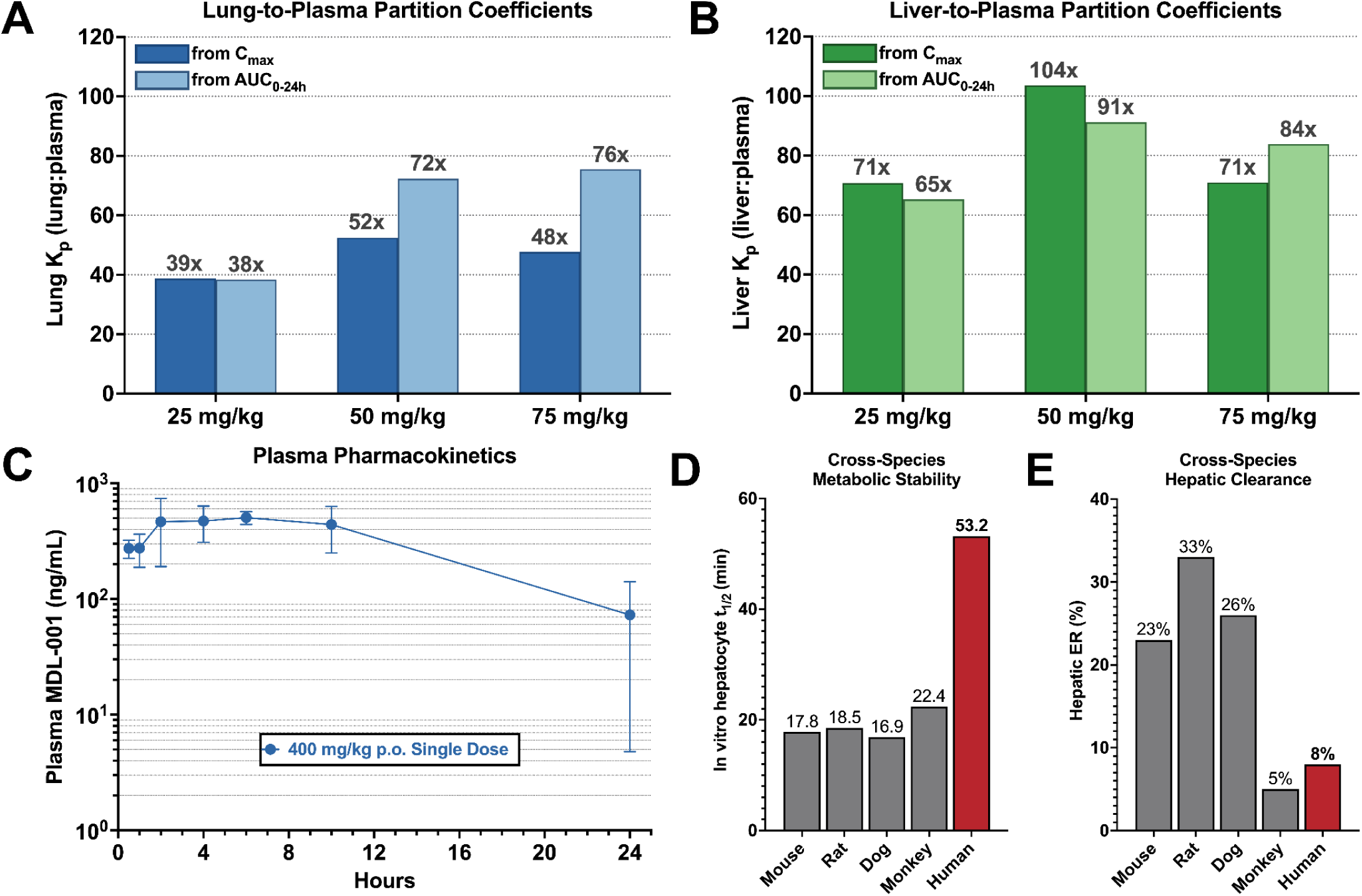
In vivo pharmacokinetics and in vitro metabolism profile of MDL-001. **(A)** Lung-to-plasma and **(B)** liver-to-plasma partition coefficients (K_p_) in CD-1 mice after single oral doses of 25, 50, or 75 mg/kg. Dark bars show K_p_ from C_max_; light bars show K_p_ from AUC0–24h. n = 5 mice per timepoint. **(C)** Plasma concentration-time profile of oral MDL-001 formulation. Mean ± SD across n = 3 male Sprague-Dawley rats per arm. **(D)** In vitro hepatocyte half-life across species. **(E)** Predicted hepatic extraction ratio across species.

The pharmacokinetics of an oral MDL-001 formulation was evaluated at 400 mg/kg (**Fig. 5C; Supplementary Table S7**). Plasma C_max_ reached 641 ng/mL.

Multi-Species Metabolism: MDL-001 Demonstrates High Human Metabolic Stability, Low Human Hepatic Clearance, and No Disproportionate Human Metabolites.

A multi-species in vitro metabolism study characterized hepatic clearance and the metabolite profile of MDL-001 in hepatocytes from mouse, rat, dog, monkey, and human (**Fig. 5D-E; Supplementary Tables S8 and S9**).

MDL-001 half-life in human hepatocytes was 53.2 minutes, exceeding mouse, rat, dog, and monkey half-lives by approximately threefold (**Fig. 5D**). The 53.2-minute human half-life established MDL-001 metabolic stability in human hepatocytes as the highest of the five species evaluated.

In vitro intrinsic clearance in human hepatocytes was 1.68 mL/min/kg, less than 10 % of total hepatic blood flow (**Supplementary Table S8**). Predicted hepatic extraction ratio in human was 8 percent and was lower than the rat value of 33 percent by approximately fourfold (**Fig. 5E; Supplementary Table S8**). The 8 percent human value classified MDL-001 as a low-clearance compound.

MDL-001 was metabolized by Phase II conjugation in all five species evaluated ( **Supplementary Table S9**). MDL-001 and four metabolites accounted for 94.1 percent of total drug-related material in the human metabolism arm. The four metabolites included two phenolic glucuronides (M10 and M11), one sulfate conjugate (M30), and one piperidine N-oxide (M1). M10 was the predominant human metabolite at 45 percent of total drug-related material. No other metabolite accounted for more than 1.5 percent of drug-related material. Every major human metabolite was detected in the toxicology species, and no metabolite unique to humans was observed. Phenolic glucuronides are rarely pharmacologically or toxicologically active [22].

#### In vitro Safety: MDL-001 is Negative in Genotoxicity Assays and Shows No Cardiac Liability

MDL-001 was evaluated in vitro in the Ames bacterial reverse-mutation assay, the micronucleus assay, and a hERG fluorescence polarization assay. MDL-001 was negative in the Ames assay, with no strain showing a 2.0-fold or greater increase in revertants. MDL-001 was also negative in the micronucleus assay, with induction remaining below the 3.0-fold positivity threshold under all conditions. MDL-001 inhibited the hERG potassium channel with an IC_50_ of 13 µM in a fluorescence polarization assay. The hERG IC_50_ exceeded the projected unbound human plasma C_max_ at the dose required to obtain a 1.0 log_10_ reduction in plasma HCV viremia by >5,000-fold (**Supplementary Table S10**).

#### In vivo Tolerability: MDL-001 Well Tolerated Across 376 Animals in Seventeen Independent Preclinical Studies

Oral and intravenous tolerability of MDL-001 was characterized across seventeen independent in vivo studies in rodents spanning single-dose and 28-day repeat-dose regimens. MDL-001 was well tolerated across 376 individual animals, with no treatment-related adverse findings by oral administration at dosing up to twice daily for 28 days (**Supplementary Table S11**).

## 3. Discussion

### MDL-001 Inhibits Positive-Sense and Negative-Sense RNA Viruses and Reverse-Transcribing DNA Viruses with Nanomolar Potency in vitro: a Direct-Acting, Broad-spectrum, Non-Nucleoside Therapeutic

No direct-acting non-nucleoside antiviral has demonstrated cross-family in vitro activity across multiple RNA virus families prior to the results reported here. MDL-001 inhibits replication of influenza A and B viruses, respiratory syncytial virus, SARS-CoV-2, endemic human coronaviruses, and hepatitis B, C, and D viruses with nanomolar potency (**Fig. 1; Supplementary Table S1**). EC_50_ values range from 27 to 263 nM across the full panel. MDL-001 inhibited viruses represent six viral families and three Baltimore Groups: positive-sense RNA viruses (Flaviviridae, Coronaviridae; Group IV), negative-sense RNA viruses (Orthomyxoviridae, Pneumoviridae; Group V), and reverse-transcribing DNA viruses (Hepadnaviridae; Group VII). This breadth of antiviral activity defies the historical one-drug-one-virus framework that has defined direct-acting antiviral development for forty years. For respiratory infection, MDL-001 establishes the in vitro foundation for a single direct-acting antiviral across influenza viruses, RSV, and SARS-CoV-2, as a potential broad-spectrum Influenza-Like Illness (ILI) therapeutic. For chronic hepatic viral disease, MDL-001 establishes the in vitro foundation for a single direct-acting antiviral across HBV, HCV, and HDV, as a potential broad-spectrum chronic hepatitis therapeutic. This antiviral activity profile also aligns MDL-001 with the broad-spectrum antiviral approach identified by NIAID as a core pandemic preparedness priority [8].

### Oral MDL-001 Establishes Monotherapy Equivalency or Superiority to Approved Standards of Care Across All Respiratory Tripledemic Viruses: a Potential broad-spectrum ILI Mono-Therapy

The absence of a broad-spectrum ILI therapy complicates treatment and compromises patient and public health. The lack of a broad-spectrum ILI treatment means patients must rearrange work, school and childcare schedules, physically travel to a healthcare site, and be administered multiple diagnostic tests prior to being prescribed and administered an antiviral treatment for an endemic respiratory mono-or co-infection. In the United States, we follow this process even though it is well understood that early therapeutic intervention in respiratory viral disease improves outcomes and saves lives. This process means patients with active infection often forgo treatment post-exposure until disease has progressed and hospitalization is required. The lack of a broad-spectrum ILI treatment means that global public health officials are as prepared to treat the next novel respiratory pandemic on Day 0 as they were to treat the previous novel pandemic on Day 0, which is to say not at all. A broad-spectrum ILI therapeutic reduces the time from exposure to treatment. It also saves hospitalizations and lives and provides the highest likelihood of a successful Day 0 treatment for the next novel respiratory pandemic.

Potential broad-spectrum ILI mono-therapies must have both broad-spectrum activity and equivalence or superiority to current standards-of-care. MDL-001 displays both properties. Oral MDL-001 establishes in vivo equivalency to or superiority over the approved standard of care for influenza and SARS-CoV-2 (**Table 1**). In a H1N1 PR8 influenza virus challenge study, oral MDL-001 was equivalent to the human equivalent clinical-dose of oseltamivir [19] for both body-weight preservation (*P = 0.81; 90%CI −1.0 to +1.9%*) and lung viral-titer reduction (90%CI −0.36 to 0.00 log_10_) on Day 5 post-infection (**Fig. 2A; Supplementary Table S2**). In a SARS-CoV-2 WA1 challenge study, oral MDL-001 reduced lung viral titers by 2.9 log_10_ (**Supplementary Table S3**), superior to the 1.4 log_10_ reduction literature reported for the human equivalent clinical-dose of nirmatrelvir (*P* = 0.023; **Table 1**) [16] and the 1.0 log_10_ reduction literature reported for the human equivalent clinical-dose of molnupiravir (*P* = 0.008; **Table 1**) [17]. MDL-001 was equivalent to the human equivalent clinical-dose of subcutaneous remdesivir [20] for body-weight preservation on Day 3 (*P = 0.47; 90%CI −3.1 to +1.3%;* **Table 1**) in the same model. In vitro, MDL-001 demonstrated an EC_50_ of 79.4 nM against respiratory syncytial virus strain A2 (**Fig. 1; Supplementary Table S1**), a 416-fold improvement over the standard of care ribavirin [18]. The data reported here demonstrate that MDL-001 is both broad-spectrum and equivalent or superior to the approved tripledemic standards of care.

**Table 1.**
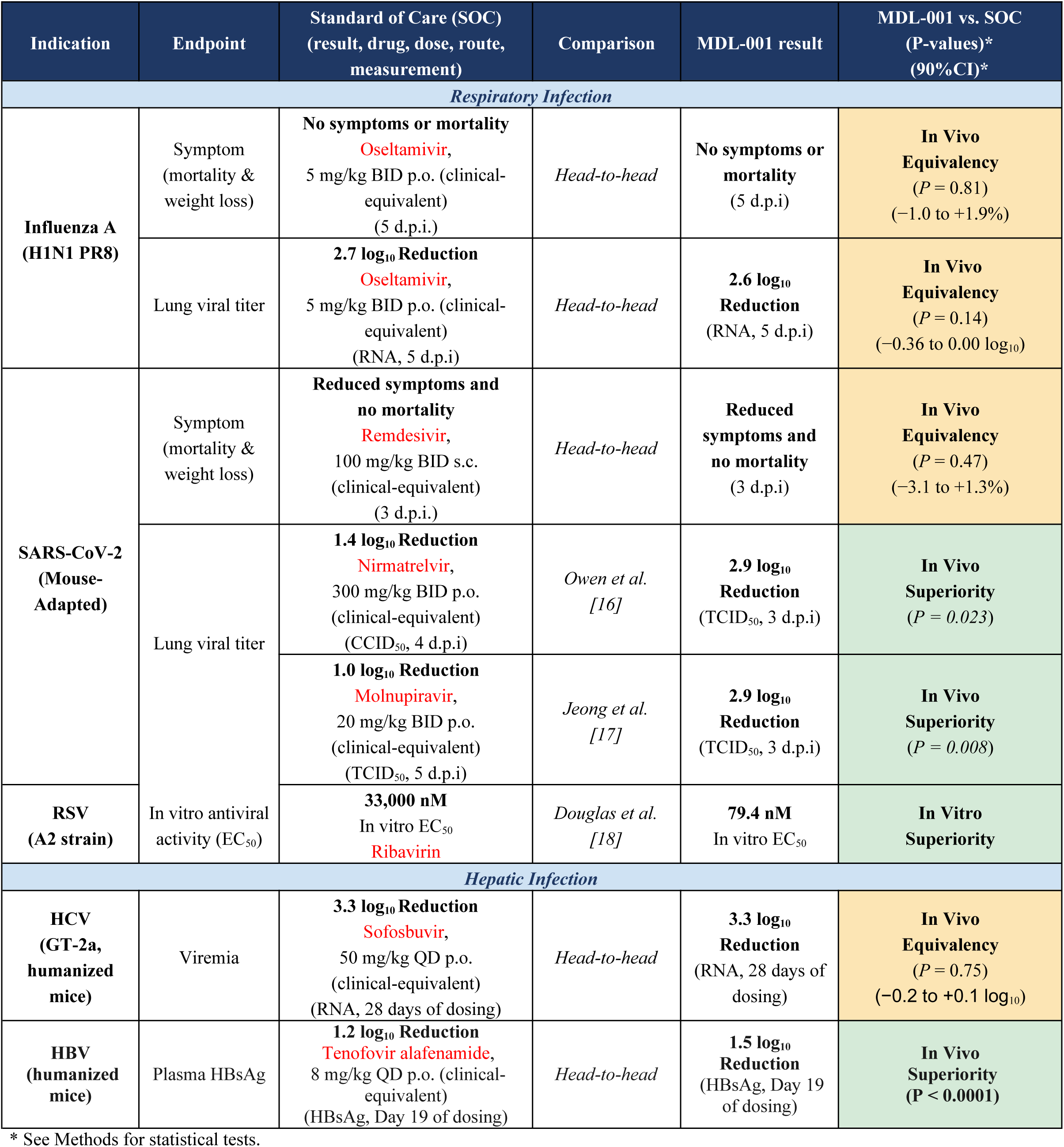
Efficacy of oral MDL-001 versus standards of care across hepatic and respiratory virus families.

Influenza viruses, SARS-CoV-2, and RSV belong to three different viral families. No approved oral therapy covers more than one. Reduction of all three by a single oral non-nucleoside small molecule directly addresses this fragmented respiratory antiviral landscape and the absence of any approved oral therapy for RSV. It also reduces the time to treatment and prepares global health officials for the next novel pandemic. Three independent prescription decisions, three diagnostic pathways, and three clinical guidelines collapse into one outpatient antiviral treatment paradigm.

### Oral MDL-001 Establishes In Vivo Equivalency to Sofosbuvir in HCV and Superiority to Tenofovir Alafenamide in HBV: a Potential Broad-Spectrum Chronic Hepatitis Therapy

The absence of a broad-spectrum chronic hepatitis therapy complicates treatment and compromises patient health and safety. An estimated 5–15 million people worldwide have both HBV and HCV infections, but fewer than 1% know their status [3–5]. Coinfection causes a multiplicative acceleration of liver disease. Patients carrying both viruses are over 100-fold more likely to develop hepatocellular carcinoma than uninfected individuals [3–4]. Furthermore, every FDA-approved HCV direct-acting antiviral (DAA) carries a black box warning for the risk of hepatitis B virus (HBV) reactivation in patients with current or prior HBV infection [6]. A broad-spectrum chronic hepatitis therapy overcomes the lack of co-infection diagnosis rates and the box warnings associated with current therapies.

Oral MDL-001 establishes in vivo equivalency to the standard of care in chronic hepatitis C and superiority to the standard of care in chronic hepatitis B. MDL-001 reduced HCV viremia by 3.3 log_10_ on Day 28 equivalent to the human equivalent dose of sofosbuvir [21] (**Fig. 3A-B; Supplementary Table S4**). MDL-001 reached equivalence with sofosbuvir at Day 21 (*P = 0.79; 90%CI 0.0 to +0.2 log_10_*), and remained equivalent at Day 24 (*P = 0.61; 90%CI −0.1 to +0.1 log_10_*) and Day 28 (*P = 0.75; 90%CI−0.2 to +0.1 log_10_*). Oral MDL-001 reduced plasma HBsAg by 2.5 log_10_ at Day 28 (**Fig. 4A-C**; **Supplementary Table S5**). MDL-001 exceeded tenofovir alafenamide at every timepoint from Day 8 through Day 21, with the largest separation at Day 19 at 1.5 against 1.2 log_10_ (P < 0.0001). Both arms converged on the assay lower limit of quantification by Day 28. The data reported here demonstrate that MDL-001 is the first single-agent therapy that addresses both HCV and HBV in vivo. MDL-001 establishes the first oral direct-acting path toward a potential broad-spectrum chronic hepatitis therapy.

### Oral MDL-001 Achieves High Lung and Liver Tissue Partitioning and Favorable Pharmacokinetics, Supporting Use as a Potential Broad-Spectrum ILI and Chronic Hepatitis Therapeutic

Tissue distribution and oral exposure are often the preclinical pharmacokinetic parameters that determine whether a lead candidate can become successful clinically. A direct-acting, broad-spectrum antiviral must achieve sufficient tissue exposure at the diverse anatomic sites those viruses occupy to be efficacious. No approved oral antiviral reaches high partitioning in both lung and liver. MDL-001 reaches both. This dual-organ exposure is the pharmacokinetic basis for the dual respiratory and hepatic indication of MDL-001.

MDL-001 target tissue/plasma ratios are higher than those of approved oral respiratory antivirals. Approved oral respiratory antivirals do not concentrate in the lung. Oseltamivir reaches a lung-to-plasma ratio near 0.5 in rats [23]. Nirmatrelvir reaches a lung-to-plasma ratio near 0.3 in hamsters [24]. The MDL-001 lung K_p_of 39 to 52 in mice exceeds the closest respiratory K_p_ by over an order of magnitude (**Fig. 5A; Supplementary Table S6**).

Approved oral hepatitis antivirals concentrate in the liver. Sofosbuvir liver/plasma ratios are nearly 30-fold in humans [25]. The MDL-001 liver K_p_ of 71 to 104 in mice sits in the upper range of what has been observed in the literature for individual components of hepatitis combination treatments (**Fig. 5B; Supplementary Table S6). Tissue accumulation** in lung and liver maps directly onto the disease anatomy of the respiratory and hepatic indications targeted by MDL-001. Pulmonary tissue exposure from oral dosing satisfies the essential tissue distribution requirements for pandemic preparedness defined by the NIAID Target Product Profiles for outpatient antivirals [7–8].

### MDL-001 Demonstrates Predominantly Phase II Metabolism Across All Species, with Metabolic Stability Highest in Humans, Supporting Standard Oral Dosing in Humans

Metabolic pathways for MDL-001 were conserved across the five species evaluated (mouse, rat, dog, monkey, human). MDL-001’s dominant metabolism was observed to be Phase II conjugation across species, generally regarded as non-toxic, detoxification biotransformations. Every major human metabolite was detected in the rodent and dog toxicology species. No human-disproportionate metabolite was observed (**Supplementary Table S9**). Equally important, no disproportionate metabolites relative to human metabolism were observed in the selected toxicology species. Conservation across toxicology species removes a common late-stage development risk and supports the toxicology species selected for IND-enabling studies.

Cross-species clearance determinations are helpful in assessing whether a candidate can be dosed orally in humans at frequencies and amounts compatible with outpatient use. In vitro intrinsic clearance in the human model is threefold lower than in rodents or dog (**Supplementary Table S8**). In vitro hepatocyte half-life in human cells exceeds rodent half-lives by threefold (**Fig. 5D**). Hepatic extraction ratio (ER_h_) in human is 8 percent (**Fig. 5E**), which classifies MDL-001 as low clearance. This profile predicts minimal hepatic first-pass metabolism in humans. Low clearance and minimal hepatic first-pass support the once- or twice-daily oral outpatient dosing frequency required for any chronic-use antiviral.

### Oral MDL-001 is Safe and Well-Tolerated In Vivo

Nucleoside analogs are the class historically proposed for broad-spectrum antiviral development [26]. The class engages host polymerases and carries chemistry-class liabilities. Ribavirin causes dose-limiting hemolytic anemia [27]. Molnupiravir is Ames-positive and carries teratogenicity concerns [26]. Remdesivir carries a hepatotoxicity warning, with transaminase elevations observed in healthy volunteers [28]. MDL-001 is a non-nucleoside that binds the Thumb-1 allosteric pocket of the viral RdRp, a site with no equivalent in human Pol II [9,29].

A broad-spectrum antiviral developed for outpatient and pandemic use must clear four standard liabilities: hERG, Ames genotoxicity, micronucleus genotoxicity, and tolerability at exposures above the minimally efficacious dose. MDL-001 clears all four. MDL-001 was well tolerated across 376 animals in seventeen independent preclinical studies spanning single-dose and 28-day repeat-dose oral regimens (**Supplementary Table S11**). MDL-001 was negative in OECD-compliant Ames and micronucleus assays. The hERG IC_50_ exceeded the projected unbound human plasma C_max_ at the minimally efficacious dose by greater than 5,000-fold (**Supplementary Table S10**). MDL-001 has documented human exposure in 79 subjects from completed Phase I studies of pipendoxifene [30–31].

### MDL-001 Meets the NIAID Broad-Spectrum Antiviral Target Product Profile and Defines a Path to Outpatient Pandemic Response, a Potential Broad-Spectrum ILI Indication and a Potential Broad-Spectrum Chronic Hepatitis Indication

No approved antiviral satisfies the NIAID strategic priority for broad-spectrum direct-acting antiviral approaches in pandemic preparedness [8]. The prevailing assumption has been that viral polymerase allosteric sites diverge too rapidly to support cross-family direct-acting antivirals. Together with our structural study [9], the data reported here refute that assumption. The Thumb-1 target family is structurally conserved and druggable across ssRNA viruses [9]. MDL-001 is the first clinical candidate reported in the literature that combines a direct-acting, non-nucleoside MoA, cross-family nanomolar activity, in vivo equivalence or superiority to Standards-of-Care, oral bioavailability, outpatient deployability, and a preclinical safety profile compatible with the attributes specified by the NIAID outpatient antiviral Target Product Profile framework [7–8].

MDL-001 is a potential first-in-class broad-spectrum therapy for broad-spectrum ILI treatment [32], broad-spectrum chronic hepatitis treatment [5], and pandemic preparedness treatment for novel viral outbreaks. Furthermore, our GALILEO AI-Drug Discovery platform and ChemPrint model have identified successive non-nucleoside chemical series targeting this family, supporting a sustained pipeline beyond MDL-001 [13]. The GALILEO platform and ChemPrint model used to discover MDL-001 have been peer-reviewed and published previously [15].

### Limitations

All drug compounds translating from in vivo proof-of-concept to human clinical development carry a general translation risk. Direct-acting antivirals (DAAs) have a general drug-class advantage in non-clinical to human translation as compared to nearly all other non-DAAs, however. DAAs, by definition, work through a target that is independent of host species and host cell line. The virus and its viral target is the same in a petri dish, in vivo and in human clinical studies, assuming the same strain is evaluated. As a result, the FDA allows cell model activity to serve as evidence of a non-clinical proof-of-concept for regulatory purposes to enter clinical trials in many instances. MDL-001 is a direct-acting antiviral.

## 4. Materials and Methods

### In Vitro Antiviral Activity Panel

The respiratory panel comprised influenza A/H1N1 wild-type, influenza A/H3N2 (MDCK), influenza B/Victoria and B/Yamagata (MDCK), respiratory syncytial virus A2 (A549), SARS-CoV-2 USA-WA1/2020 and variants of concern Alpha, Beta, Delta, and Omicron (HeLa-ACE2), mouse-adapted SARS-CoV-2 WA1 (HeLa-ACE2), human alphacoronavirus 229E (MRC-5), and human betacoronavirus OC43 (A549-ACE2). The hepatic panel comprised hepatitis C virus genotypes 1b and 2a in subgenomic replicon systems [33], hepatitis D virus genotype I in hNTCP-expressing Huh7 cells with HBV co-infection [34], and hepatitis B virus in the HepAD38 inducible cell system [35]. Complete assay parameters for each virus-cell combination are provided in **Supplementary Table S1**.

Antiviral activity was measured in cell-based infection assays using cytopathic effect, immunofluorescence, or viral yield reduction readouts as appropriate for each virus. Concentration-response curves (4 to 10 concentrations) were generated from technical replicates and fit by four-parameter logistic regression to estimate EC_50_ and EC_90_ values. Cytotoxicity controls were run in parallel through either MTT or Cell-Titer-Glo. EC_50_ and EC_90_ values are plotted in **Fig. 1**.

### FBS Protein Binding

Protein binding of MDL-001 was determined by rapid equilibrium dialysis (RED) in fetal bovine serum (FBS). MDL-001 (1 µM) was incubated with 2% and 10% FBS under both heat-inactivated and non-heat-inactivated conditions. The unbound fraction (F_ub_) at 10% FBS was 5.0% (heat-inactivated) and 4.7% (non-heat-inactivated). The unbound fraction at 2% FBS was 25.0% (heat-inactivated) and 24.4% (non-heat-inactivated). Mean recovery ranged from 91.2% to 112.4% across all conditions. These values were used to calculate free-fraction antiviral EC_50_ and EC_90_ values reported in **Fig. 1** and **Supplementary Table S1.**

### Influenza A In Vivo Challenge

Balb/c mice were anesthetized and inoculated intranasally with 1.0 × 10^5^ PFU of influenza A/Puerto Rico/8/34 (PR8) luciferase reporter virus, approximately 1,900-fold the reported murine intranasal 50% lethal dose for A/PR/8/34 [36]. Animals were randomized to three treatment groups with n = 3 per group: vehicle (formulation blank), MDL-001 at 400 mg/kg by oral gavage twice daily, and oseltamivir at 5 mg/kg by oral gavage twice daily, the mouse equivalent of the approved human clinical dose [19]. The first dose of all treatments was administered 24 hours before infection and dosing continued through Day 5. All work was conducted under BSL-2 conditions under IACUC-approved protocols. The two experimental arms and the vehicle arm were compared to evaluate the superiority, equivalence, or inferiority of MDL-001 to oseltamivir in reducing symptomatic disease progression and lung viral titer at Day 5. Symptomatic disease progression was measured as body-weight loss and mortality.

Body weight was recorded twice daily through Day 5 post-infection. Surviving mice were euthanized on Day 5 post-infection. Lungs were harvested, and viral burden was quantified by RT-qPCR for H1N1 PR8 RNA expressed as copies per milligram of lung tissue. Body weight time courses were analyzed by two-way ANOVA as percent-baseline values with comparisons to vehicle. Day 5 lung titers were analyzed by two-way ANOVA on log_10_-transformed values with comparisons to vehicle. Significance thresholds were *P < 0.05, **P < 0.01, ***P < 0.001, and ****P < 0.0001. Body weight time courses are shown in **Fig. 2A**. Day 5 lung burden is tabulated in **Supplementary Table S2**.

### SARS-CoV-2 In Vivo Challenge

Female 10-week-old 129S1/SvImJ mice (Jackson Laboratory strain 002448) were anesthetized and inoculated intranasally with 2.5 x 10^4^ PFU of mouse-adapted SARS-CoV-2 WA1/2020. Animals were randomized to treatment groups with n = 9 per group. MDL-001 was administered in 0.5% methylcellulose by oral gavage at 250 mg/kg twice daily for 3 days. Remdesivir was administered subcutaneously at 100 mg/kg twice daily, the human equivalent dose [20], for 3 days. The mouse-adapted SARS-CoV-2 used here was derived by serial passage of USA-WA1/2020 in mouse lungs and has been published extensively [37–38]. Vehicle-treated animals received 0.5% methylcellulose by oral gavage. The first dose of all treatments was administered 1 hour before intranasal infection. Body weight was measured daily through Day 3 post-infection. Body weight time courses were analyzed by two-way ANOVA as percent-baseline values with comparisons to vehicle. Significance thresholds were *P < 0.05, **P < 0.01, ***P < 0.001, and ****P < 0.0001. Body weight time courses are shown in **Fig. 2B**. All work was conducted under BSL-3 conditions under IACUC-approved protocols. The experimental arms and the vehicle arm were compared to evaluate the superiority, equivalence, or inferiority of MDL-001 to remdesivir in reducing body-weight loss at Day 3.

Female 10-week-old 129S1/SvImJ mice (Jackson Laboratory strain 002448) were anesthetized and inoculated intranasally with 2.5 x 10^4^ PFU of mouse-adapted SARS-CoV-2 WA1/2020. Animals were randomized to treatment groups with n = 9 per group. MDL-001 was administered in 0.5% methylcellulose by oral gavage at 250 mg/kg twice daily for 3 days. The mouse-adapted SARS-CoV-2 used here was derived by serial passage of USA-WA1/2020 in mouse lungs and has been published extensively [37–38]. On Day 3, mice were euthanized, lungs were harvested, homogenized in PBS with silica glass beads, and stored at –80°C. Infectious virus was quantified by limiting-dilution titration on Vero-TMPRSS2 cells. Cells were seeded at 20,000 cells per well in 96-well plates. Serial dilutions of clarified lung homogenate were applied. Cytopathic effect was scored at 5 days by crystal violet staining. TCID_50_ values were calculated by the Reed-Muench method and expressed as TCID_50_/mL. Day 3 lung titers were analyzed by two-way ANOVA on log_10_-transformed values with comparisons to vehicle. Significance thresholds were *P < 0.05, **P < 0.01, ***P < 0.001, and ****P < 0.0001. Day 3 lung titers are tabulated in **Supplementary Table S3**. All work was conducted under BSL-3 conditions under IACUC-approved protocols. MDL-001 and the literature-reported clinical equivalent doses of nirmatrelvir and molnupiravir were compared to evaluate the superiority, equivalence, or inferiority of MDL-001 to these literature reported values.

### HCV In Vivo Challenge

Transgenic MUP-uPA-SCID/Beige mice were engrafted intrasplenically with 1 x 10^7^ primary human hepatocytes. Successful humanization was confirmed by serum human albumin greater than 300 µg/mL before randomization. Human albumin was quantified by ELISA at baseline and at terminal harvest to confirm that the human hepatocyte graft remained intact for the duration of dosing. This humanized-liver mouse model supports productive infection with HCV as previously described [39–40]. Animals were randomized to treatment arms with n = 6 per group. Mice were infected intravenously on Day −2 with 1 x 10^6^ IU HCV JFH1/J6.

Treatments were administered by oral gavage. Study arms included vehicle, the human equivalent dose of sofosbuvir 50 mg/kg once daily [21], and MDL-001 400 mg/kg twice daily for 28 days. Blood was collected on Day 0 (baseline) and every 3 days thereafter through Day 28. Plasma HCV RNA copies were quantified by TaqMan RT-qPCR. The experimental arms and the vehicle arm were compared to evaluate the superiority, equivalence, or inferiority of MDL-001 to sofosbuvir in reducing HCV RNA over 28 days.

Plasma viral loads are reported as copies per milliliter and as a percentage of the contemporaneous vehicle mean. Log_10_ reductions were calculated as log_10_(mean vehicle / mean treatment). Statistical comparisons were performed using two-way ANOVA (treatment × time) on log_10_-transformed values, with vehicle as the reference. Significance thresholds were *P < 0.05, **P < 0.01, ***P < 0.001, and ****P < 0.0001. Plasma viral load data are plotted in **Fig. 3A-B**. Day-by-day log_10_ reductions are tabulated in **Supplementary Table S4**.

### HBV In Vivo Challenge

Transgenic MUP-uPA-SCID/Beige mice were engrafted intrasplenically with 1 x 10^7^ primary human hepatocytes. Successful humanization was confirmed by serum human albumin greater than 300 µg/mL before randomization. Human albumin was quantified by ELISA at baseline and at terminal harvest to confirm that the human hepatocyte graft remained intact for the duration of dosing. This humanized-liver mouse model supports productive infection with HBV as previously described [39]. Animals were randomized to treatment arms with n = 6 per group. Mice were infected intravenously on Day −2 with 1 x 10^7^ genome equivalents of HBV (AD38 strain).

Treatments were administered by oral gavage. Study arms included vehicle, tenofovir alafenamide 8 mg/kg once daily, and MDL-001 400 mg/kg twice daily for 28 days. Blood was collected on Day 0 (baseline), daily on Days 1 through 10, and on Days 13, 16, 19, 21, 24, 27, and 28. Plasma HBsAg was quantified by ELISA. The lower limit of quantification was 1 ng/mL. The two experimental arms and the vehicle arm were compared to evaluate the superiority, equivalence, or inferiority of MDL-001 to tenofovir alafenamide in reducing plasma HBsAg over 28 days.

Plasma HBsAg is reported as nanograms per milliliter and as a percentage of the contemporaneous vehicle mean. Log10 reductions were calculated as log10(mean vehicle / mean treatment). Significance versus vehicle was tested by two-way ANOVA (treatment × time) on log10-transformed values, with vehicle as the reference. Significance thresholds were *P < 0.05, **P < 0.01, ***P < 0.001, and ****P < 0.0001. The MDL-001 versus tenofovir alafenamide comparison was performed on log10-transformed per-animal values by two-sided unpaired Welch’s unequal-variance t-test. Values at or below the lower limit of quantification were censored at the limit before transformation, so reductions on days when an arm reached the limit are lower bounds. Plasma HBsAg data are plotted in **Fig. 4A-C**. Day-by-day log10 reductions are tabulated in **Supplementary Table S5**.

### Standard of Care Comparison Statistics

Calculation of averages and confidence intervals: Each treatment group’s mean and two-sided 95% confidence interval were calculated from the per-animal values using Student’s t distribution for each treatment arm at each timepoint.

Calculation of Superiority: Active-comparator P values in Table 1 were calculated with two-sided unpaired Welch’s unequal-variance t-tests to test superiority. Welch-Satterthwaite degrees of freedom were used for all Welch tests. Superiority is determined if the calculated p value <0.05 and the CI 95% of the difference does not cross 0.

Calculation of Equivalence: Where superiority was not achieved, equivalence to the standard of care comparator was assessed by two one-sided tests (TOST) [41]. This procedure follows the approach recommended by the U.S. Food and Drug Administration for establishing equivalence [42]. The equivalence limits were ±0.5 log10 for viral-titer and viral-load reductions and ±5% of baseline for body weight. For each endpoint the difference between MDL-001 and the comparator was summarized as a 90% confidence interval, and equivalence was concluded when that interval fell entirely within the limit.

Influenza body weights were expressed as percent of each animal’s Day-0 baseline. AM and PM readings were averaged within animal to generate one daily value. Body-weight preservation was calculated as active-treated animal daily percent-baseline body weight minus the contemporaneous vehicle daily mean. MDL-001 and oseltamivir distributions were compared on Days 1 through 5. Influenza lung viral RNA was analyzed as animal-level absolute reductions from vehicle. Absolute reduction was calculated as the contemporaneous vehicle arithmetic mean RNA copies/mg lung minus the individual active-treated animal value. MDL-001 and oseltamivir distributions did not differ detectably at Day 5. These statistics are reported in **Supplementary Table S2**.

SARS-CoV-2 body weights were expressed as percent of each animal’s Day-0 baseline. Body-weight preservation was calculated as active-treated animal percent-baseline body weight minus the contemporaneous vehicle mean. MDL-001 and remdesivir distributions were compared on Days 1 through 3. SARS-CoV-2 literature-comparator *P* values were calculated with two-sided Welch tests on absolute log_10_ viral-titer reduction summary statistics. The MDL-001 summary statistic was calculated from raw Day 3 animal-level lung viral-titer reductions relative to the contemporaneous vehicle geometric mean. The MDL-001 values were mean = 2.9 log_10_, SD = 1.605, and n = 9. The nirmatrelvir summary statistic was extracted from Owen et al. [16]. 300 mg/kg BID nirmatrelvir yielded a 1.4 log_10_ reduction (CCID_50_, 4 d.p.i) with SD of 0.648. The molnupiravir summary statistic was extracted from Jeong et al. [17]. 20 mg/kg BID molnupiravir yielded a 1.0 log_10_ reduction (TCID_50_, 5 d.p.i), with an SD of 0.508. These statistics are reported in **Supplementary Table S3**.

HCV plasma viremia was analyzed as animal-level absolute reduction from vehicle. Absolute reduction was calculated as the contemporaneous vehicle arithmetic mean HCV RNA copies/mL minus the individual active-treated animal value. MDL-001 and sofosbuvir distributions were compared on Days 21, 24, and 28.

HBV plasma HBsAg was analyzed as animal-level absolute reduction from vehicle. Absolute reduction was calculated as the contemporaneous vehicle arithmetic mean HBsAg ng/mL minus the individual active-treated animal value. MDL-001 and tenofovir alafenamide distributions were compared on Days 8, 9, 10, 13, 16, 19, 21, and 24.

### In Vivo Pharmacokinetics and Tissue Distribution

CD-1 mice received single oral doses of 25, 50, or 75 mg/kg MDL-001 formulated as an oral suspension. Plasma, lung, and liver were collected at 1, 2, 4, 8, 16, and 24 h post-dose. Five mice contributed samples at each timepoint, for a total of 30 mice per dose level and 90 mice across the study. MDL-001 concentrations in plasma and tissues were quantified by validated LC-MS/MS in accordance with FDA bioanalytical guidance [43]. Tissues were homogenized prior to extraction. Verapamil (200 ng/mL) served as an internal standard. Lower limits of quantitation were 1 ng/mL for plasma and 50 ng/g for lung and liver. Bioanalysis was performed at Frontage Laboratories.

Tissue-to-plasma partition coefficients (K_p_ = C_tissue_/C_plasma_) were calculated as the ratio of peak tissue concentration to matched peak plasma concentration, and as the ratio of tissue AUC0–24h to plasma AUC0–24h. Per-dose K_p_ values across the 25, 50, and 75 mg/kg dose levels are reported in **Fig. 5A-B**. Calculated oral pharmacokinetic parameters are listed in **Supplementary Table S6**.

### In Vivo Plasma Pharmacokinetics

Plasma exposure of MDL-001 was determined in the rat using an optimized oral solution formulation at Frontage Laboratories. Male Sprague-Dawley rats received a single oral dose at 400 mg/kg (n = 3). Plasma was sampled at 0.5, 1, 2, 4, 6, 10, and 24 h post-dose. MDL-001 was quantified by a qualified LC-MS/MS method. C_max_, C_24h_, t_1/2_, AUC_last_, and AUC_inf_ were derived by non-compartmental analysis. Plasma concentration-time profiles are plotted in **Fig. 5C**. Calculated pharmacokinetic parameters are listed in **Supplementary Table S7**.

### Hepatocyte Stability and Hepatic Clearance Prediction

Metabolic stability was measured in pooled cryopreserved hepatocytes from mouse, rat, dog, cynomolgus monkey, and human at 1 µM MDL-001 and 1 x 10^6^ cells/mL. Compound disappearance was quantified by UPLC-MS, and half-life and intrinsic clearance were derived from first-order kinetics. Predicted in vivo hepatic clearance (mL/min/kg) was calculated using the well-stirred model [44–45] with species-specific hepatocellularity, liver blood flow, and measured plasma fraction unbound. Blood-to-plasma ratio was assumed to be 1. Hepatocyte half-life and predicted hepatic extraction ratio across species are plotted in **Fig. 5D-E**. Plasma protein binding across species was determined by rapid equilibrium dialysis (**Supplementary Tables S8 and S9**).

### Genetic Toxicology (Ames & MNT assays)

Mutagenicity was assessed using the Ames MPF microplate format [46] in accordance with OECD Test Guideline 471 [47]. Salmonella typhimurium strains TA98, TA100, TA1535, and TA1537 and Escherichia coli WP2 uvrA[pKM101] were tested at six concentrations (0.06 to 62.5 µg/mL) with and without rat liver S9 metabolic activation. Positive controls were strain-appropriate. A 2.0-fold or greater increase in revertants with dose-response was the positivity criterion.

Clastogenicity was assessed in CHO-K1 cells according to OECD Test Guideline 487 [48]. Cells were exposed to an 11-point concentration series spanning 0.03 to 31.3 µM with S9 (4 h treatment plus 20 h recovery) or without S9 (24 h continuous exposure). Micronuclei were quantified by flow cytometry with dual DNA staining. Cyclophosphamide (+S9) and vinblastine (-S9) served as positive controls. Induction less than 3.0-fold was considered negative.

### Cardiac Safety (hERG assay)

Binding to the hERG potassium channel was measured using the Predictor hERG fluorescence polarization assay (Thermo Fisher Scientific) per the manufacturer’s protocol. A 16-point dilution series of MDL-001 (0.007 nM to 100 µM) was incubated with hERG membrane preparations and fluorescent tracer for 4 h at room temperature. Fluorescence polarization was read on a plate reader, and IC_50_ was determined by fitting percent inhibition against concentration with a four-parameter logistic model. E-4031 and atenolol served as positive and negative controls, respectively. The Predictor assay buffer contains no plasma protein. Compound concentrations of 33 µM and 100 µM showed non-specific assay interference and were excluded from the fit. The cardiac safety margin was calculated as the ratio of the hERG IC_50_ to the projected unbound human plasma C_max_ at the minimally efficacious dose, using species-specific plasma protein binding determined by rapid equilibrium dialysis (**Supplementary Table S10**). The minimum efficacious dose is defined as the dose required to achieve a 1-log reduction in HCV viremia at Day 10 in mice: currently 62.5 mg/kg.

### Preclinical Tolerability

Tolerability data were compiled from seventeen independent in vivo experiments conducted at multiple institutions, including efficacy, pharmacokinetic, and dedicated safety observations. Animals were monitored for clinical signs, body weight changes, and mortality. Studies included dosing up to 28-day BID administered by oral gavage. The complete inventory of studies, durations, animal numbers, and routes is provided in **Supplementary Table S11**.

## Acknowledgments

The authors thank Dr. George Nicola, co-inventor with Dr. Daniel Haders, of MDL-001. The authors thank Dr. Tushar Menon and Dr. Launa Aspeslet for expert consultation and detailed review of this manuscript. The authors thank Dr. Luis Martinez-Sobrido for supplying the constructs of influenza virus.

## Supplementary Materials

The following supporting information is available with this preprint. Table S1: In vitro antiviral activity of MDL-001 by virus family, virus, and assay system; Table S2: Lung log10 viral reduction from vehicle (Influenza Study); Table S3: Lung log10 viral reduction from vehicle (SARS-CoV-2 Study); Table S4: Plasma log10 viral reduction from vehicle for each day (HCV Study); Table S5: Plasma HBsAg log10 reduction from vehicle for each day (HBV Study); Table S6: Oral pharmacokinetics of MDL-001 in mouse plasma, lung, and liver after single oral doses; Table S7: Rat plasma pharmacokinetics of oral MDL-001 formulation; Table S8: Cross-species in vitro intrinsic clearance of MDL-001; Table S9: Metabolite profile for MDL-001 across species in cryopreserved hepatocytes; Table S10: Calculation of hERG safety margin from in vitro IC50 and projected unbound human plasma Cmax; Table S11: Summary of duration, animal, and route of administration for the tolerability studies of MDL-001 in mice and rats.

## Author Contributions

Conceptualization, D.H., T.U. and V.W.; Methodology, D.H.; Investigation, P.G., A.G., M.B., B.L.M., R.R., M.L.R., H.v.B., E.M.S., V.S., A.G.-S. and K.M.W.; Formal analysis, D.H., T.U., V.W., S.R. and D.S.; Writing-original draft, V.W., T.U. and D.H.; Writing - review & editing, all authors; Visualization, V.W., D.H.; Supervision, A.G.-S., K.M.W., P.G., D.S. and D.H.; Project administration, D.H. All authors have read and agreed to the published version of the manuscript.

## Institutional Review Board Statement

All animal procedures followed protocols approved by the institutional animal care and use committees at each study site and complied with the Guide for the Care and Use of Laboratory Animals (National Research Council, 2011). The influenza A challenge studies were approved by the Institutional Animal Care and Use Committee of The Scripps Research Institute under protocol 20-0007 (approved 8 May 2026). The HCV, HBV, and mouse pharmacokinetic studies were approved under Scripps Research protocol 11-0015 (approved 27 February 2026). The rat pharmacokinetic study was approved at Frontage Laboratories under protocols 2025-1005B and 2025-1001C (both approved 21 May 2026), and the rats were obtained from Charles River Laboratories. The SARS-CoV-2 challenge studies were approved by the Institutional Animal Care and Use Committee of the Icahn School of Medicine at Mount Sinai under protocol PROTO202100056 (approved 20 May 2021).

## Informed Consent Statement

Not applicable.

## Data Availability Statement

The original contributions presented in this study are included in the article/Supplementary Materials. Further inquiries can be directed to the corresponding author(s).

## References

1. Luong QXT, Hoang PT, Ho PT, Ayun RQ, Lee TK, Lee S. 2025. Potential broad-spectrum antiviral agents: a key arsenal against newly emerging and reemerging respiratory RNA viruses. Int J Mol Sci 26:1481. 10.3390/ijms26041481

2. Rossignol JF. 2022. Rethinking methods used to evaluate effectiveness of therapeutics for COVID-19 and other viral respiratory illnesses. Future Virol 17:67–69. 10.2217/fvl-2021-0209

3. Mavilia MG, Wu GY. 2018. HBV-HCV coinfection: viral interactions, management, and viral reactivation. J Clin Transl Hepatol 6:296–305. 10.14218/JCTH.2018.00016

4. Maqsood Q, Sumrin A, Iqbal M, Younas S, Hussain N, Mahnoor M, Wajid A. 2023. Hepatitis C virus/hepatitis B virus coinfection: current prospectives. Antivir Ther 28:13596535231189643. 10.1177/13596535231189643

5. World Health Organization. 2024. Global hepatitis report 2024. World Health Organization, Geneva. https://www.who.int/publications/i/item/9789240091672

6. Bersoff-Matcha SJ, Cao K, Jason M, Ajao A, Jones SC, Meyer T, Brinker A. 2017. Hepatitis B virus reactivation associated with direct-acting antiviral therapy for chronic hepatitis C virus: a review of cases reported to the U.S. Food and Drug Administration adverse event reporting system. Ann Intern Med 166:792–798. 10.7326/M17-0377

7. National Institute of Allergy and Infectious Diseases. 2023. Target product profiles for antivirals. National Institutes of Health, Bethesda, MD. https://www.niaid.nih.gov/research/target-product-profiles-antivirals

8. National Institute of Allergy and Infectious Diseases. 2022. NIAID pandemic preparedness plan. National Institutes of Health, Bethesda, MD. https://www.niaid.nih.gov/research/pandemic-preparedness

9. Woods V, Umansky T, Russell SM, Gallay P, Smith D, Haders D. 2024. Discovery of RdRp Thumb-1 as a novel broad-spectrum antiviral family of targets and MDL-001 as a potent broad-spectrum inhibitor thereof. bioRxiv. 10.1101/2024.03.29.587401

10. Gentile I, Zappulo E, Buonomo AR, Maraolo AE, Borgia G. 2015. Beclabuvir for the treatment of hepatitis C. Expert Opin Investig Drugs 24:1111–1121. 10.1517/13543784.2015.1059820

11. Gentles RG. 2019. Discovery of beclabuvir: a potent allosteric inhibitor of the hepatitis C virus polymerase. Top Med Chem 31:193–228. 10.1007/7355_2018_38

12. Umansky T, Woods V, Russell SM, Smith D, Haders D. 2024. ChemPrint: an AI-driven framework for enhanced drug discovery. bioRxiv. 10.1101/2024.03.22.586314

13. Umansky T, Woods V, Russell SM, Garvey DS, Smith D, Haders D. 2025. GALILEO generatively expands chemical space and achieves one-shot identification of a library of novel, specific, next generation broad-spectrum antiviral compounds at high hit rates. bioRxiv. 10.1101/2025.01.17.633620

14. Umansky TJ, Woods VA, Russell SM, Haders DJ. 2026. AmesNet: a task-conditioned deep learning model with enhanced sensitivity and generalization in Ames mutagenicity prediction. Chem Res Toxicol. 10.1021/acs.chemrestox.6c00082

15. Haders D, Nicola G. 2026. Methods and compositions for treating RNA viral infections. US patent 12,551,475.

16. Owen DR, Allerton CMN, Anderson AS, Aschenbrenner L, Avery M, Berritt S, Boras B, Cardin RD, Carlo A, Coffman KJ, Dantonio A, Di L, Eng H, Ferre R, Gajiwala KS, Gibson SA, Greasley SE, Hurst BL, Kadar EP, Kalgutkar AS, Lee JC, Lee J, Liu W, Mason SW, Noell S, Novak JJ, Obach RS, Ogilvie K, Patel NC, Pettersson M, Rai DK, Reese MR, Sammons MF, Sathish JG, Singh RSP, Steppan CM, Stewart AE, Tuttle JB, Updyke L, Verhoest PR, Wei L, Yang Q, Zhu Y. 2021. An oral SARS-CoV-2 Mpro inhibitor clinical candidate for the treatment of COVID-19. Science 374:1586–1593. 10.1126/science.abl4784

17. Jeong JH, Chokkakula S, Min SC, Kim BK, Choi WS, Oh S, Yun YS, Kang DH, Lee OJ, Kim EG, Choi JH, Lee JY, Choi YK, Baek YH, Song MS. 2022. Combination therapy with nirmatrelvir and molnupiravir improves the survival of SARS-CoV-2 infected mice. Antiviral Res 208:105430. 10.1016/j.antiviral.2022.105430

18. Douglas JL, Panis ML, Ho E, Lin K-Y, Krawczyk SH, Grant DM, Cai R, Swaminathan S, Cihlar T. 2003. Inhibition of respiratory syncytial virus fusion by the small molecule VP-14637 via specific interactions with F protein. J Virol 77:5054–5064. 10.1128/jvi.77.9.5054-5064.2003

19. Döhrmann S, Levin J, Cole JN, Borchardt A, Amundson K, Almaguer A, Abelovski E, Grewal R, Zuill D, Dedeic N, Hough G, Fortier J, Donatelli J, Lam T, Chen ZY, Jiang W, Haussener T, Noncovich A, Balkovec JM, Bensen DC, Ong V, Brady TP, Locke JB, Flanagan S, Hughes RM, Stein JL, Tari LW. 2025. Drug-Fc conjugate CD388 targets influenza virus neuraminidase and is broadly protective in mice. Nat Microbiol 10:912–926. 10.1038/s41564-025-01955-3

20. Pruijssers AJ, George AS, Schäfer A, Leist SR, Gralinski LE, Dinnon KH III, Yount BL, Agostini ML, Stevens LJ, Chappell JD, Lu X, Hughes TM, Gully K, Martinez DR, Brown AJ, Graham RL, Perry JK, Du Pont V, Pitts J, Ma B, Babusis D, Murakami E, Feng JY, Bilello JP, Porter DP, Cihlar T, Baric RS, Denison MR, Sheahan TP. 2020. Remdesivir inhibits SARS-CoV-2 in human lung cells and chimeric SARS-CoV expressing the SARS-CoV-2 RNA polymerase in mice. Cell Rep 32:107940. 10.1016/j.celrep.2020.107940

21. Mesci P, Macia A, Moore SM, Shiryaev SA, Pinto A, Huang CT, Tejwani L, Fernandes IR, Suarez NA, Kolar MJ, Montefusco S, Rosenberg SC, Herai RH, Cugola FR, Russo FB, Sheets N, Saghatelian A, Shresta S, Momper JD, Siqueira-Neto JL, Corbett KD, Beltrão-Braga PCB, Terskikh AV, Muotri AR. 2018. Blocking Zika virus vertical transmission. Sci Rep 8:1218. 10.1038/s41598-018-19526-4

22. Stachulski AV, Meng X. 2013. Glucuronides from metabolites to medicines: a survey of the in vivo generation, chemical synthesis and properties of glucuronides. Nat Prod Rep 30:806–848. 10.1039/C3NP70003H

23. Gao G, Law FCP, Wong RNS, Mak NK, Yang MSM. 2019. A physiologically-based pharmacokinetic model of oseltamivir phosphate and its carboxylate metabolite for rats and humans. ADMET DMPK 7:22–43. 10.5599/admet.628

24. Abdelnabi R, Jochmans D, Donckers K, Trüeb B, Ebert N, Weynand B, Thiel V, Neyts J. 2023. Nirmatrelvir-resistant SARS-CoV-2 is efficiently transmitted in female Syrian hamsters and retains partial susceptibility to treatment. Nat Commun 14:2124. 10.1038/s41467-023-37773-6

25. Babusis D, Curry MP, Kirby B, Park Y, Murakami E, Wang T, Mathias A, Afdhal N, McHutchison JG, Ray AS. 2018. Sofosbuvir and ribavirin liver pharmacokinetics in patients infected with hepatitis C virus. Antimicrob Agents Chemother 62:e02587–17. 10.1128/AAC.02587-17

26. Karim M, Lo CW, Einav S. 2023. Preparing for the next viral threat with broad-spectrum antivirals. J Clin Invest 133:e170236. 10.1172/JCI170236

27. Russmann S, Grattagliano I, Portincasa P, Palmieri VO, Palasciano G. 2006. Ribavirin-induced anemia: mechanisms, risk factors and related targets for future research. Curr Med Chem 13:3351–3357. 10.2174/092986706778773059

28. Humeniuk R, Mathias A, Cao H, Osinusi A, Shen G, Chng E, Ling J, Vu A, German P. 2020. Safety, tolerability, and pharmacokinetics of remdesivir, an antiviral for treatment of COVID-19, in healthy subjects. Clin Transl Sci 13:896–906. 10.1111/cts.12840

29. Wu J, Wang X, Liu Q, Lu G, Gong P. 2023. Structural basis of transition from initiation to elongation in de novo viral RNA-dependent RNA polymerases. Proc Natl Acad Sci U S A 120:e2211425120. 10.1073/pnas.2211425120

30. Gandhi T, Krishnamurthy R, Patel S, Feldman D, MacRae L, Trammel DK, van Haarlem T, Cotreau MM, Galbreath E. 2000. Safety and pharmacokinetic evaluation of ERA-923 in healthy postmenopausal women in two double masked phase I trials. Ann Oncol 11:41.

31. Cotreau MM, Stonis L, Dykstra KH, Gandhi T, Gutierrez M, Xu J, Park Y, Burghart PH, Schwertschlag US. 2002. Multiple-dose, safety, pharmacokinetics, and pharmacodynamics of a new selective estrogen receptor modulator, ERA-923, in healthy postmenopausal women. J Clin Pharmacol 42:157–165. 10.1177/00912700222011193

32. Centers for Disease Control and Prevention. 2022. Increased respiratory virus activity, especially among children, early in the 2022-2023 fall and winter. CDC Health Alert Network, November 4, 2022. https://emergency.cdc.gov/han/2022/han00479.asp

33. Lohmann V, Korner F, Koch JO, Herian U, Theilmann L, Bartenschlager R. 1999. Replication of subgenomic hepatitis C virus RNAs in a hepatoma cell line. Science 285:110–113. 10.1126/science.285.5424.110

34. Ni Y, Lempp FA, Mehrle S, Nkongolo S, Kaufman C, Falth M, Stindt J, Koniger C, Nassal M, Kubitz R, Sultmann H, Urban S. 2014. Hepatitis B and D viruses exploit sodium taurocholate co-transporting polypeptide for species-specific entry into hepatocytes. Gastroenterology 146:1070–1083. 10.1053/j.gastro.2013.12.024

35. Ladner SK, Otto MJ, Barker CS, Zaifert K, Wang GH, Guo JT, Seeger C, King RW. 1997. Inducible expression of human hepatitis B virus (HBV) in stably transfected hepatoblastoma cells: a novel system for screening potential inhibitors of HBV replication. Antimicrob Agents Chemother 41:1715–1720. 10.1128/AAC.41.8.1715

36. Bowen LE, Rivers K, Trombley JE, Bohannon JK, Li SX, Boydston JA, Eichelberger MC. 2012. Development of a murine nose-only inhalation model of influenza: comparison of disease caused by instilled and inhaled A/PR/8/34. Front Cell Infect Microbiol 2:74. 10.3389/fcimb.2012.00074

37. Rathnasinghe R, Jangra S, Cupic A, Martínez-Romero C, Mulder LCF, Kehrer T, Yildiz S, Choi A, Mena I, De Vrieze J, Aslam S, Stadlbauer D, Meekins DA, McDowell CD, Balaraman V, Richt JA, De Geest BG, Miorin L, Krammer F, Simon V, García-Sastre A, Schotsaert M. 2022. Characterization of SARS-CoV-2 Spike mutations important for infection of mice and escape from human immune sera. Nat Commun 13:3921. 10.1038/s41467-022-30763-0

38. Rosales R, McGovern BL, Rodriguez ML, Leiva-Rebollo R, Diaz-Tapia R, Benjamin J, Rai DK, Cardin RD, Anderson AS, PSP study group, Sordillo EM, van Bakel H, Simon V, García-Sastre A, White KM. 2024. Nirmatrelvir and molnupiravir maintain potent in vitro and in vivo antiviral activity against circulating SARS-CoV-2 omicron subvariants. Antiviral Res 230:105970. 10.1016/j.antiviral.2024.105970

39. Tesfaye A, Stift J, Maric D, Cui Q, Dienes HP, Feinstone SM. 2013. Chimeric mouse model for the infection of hepatitis B and C viruses. PLoS One 8:e77298. 10.1371/journal.pone.0077298

40. Bobardt M, Ramirez CM, Baum MM, Ure D, Foster R, Gallay PA. 2021. The combination of the NS5A and cyclophilin inhibitors results in an additive anti-HCV inhibition in humanized mice without development of resistance. PLoS One 16:e0251934. 10.1371/journal.pone.0251934

41. Schuirmann DJ. 1987. A comparison of the two one-sided tests procedure and the power approach for assessing the equivalence of average bioavailability. J Pharmacokinet Biopharm 15:657–680. 10.1007/BF01068419

42. U.S. Food and Drug Administration. 2026. Statistical approaches to establishing bioequivalence guidance for industry. Center for Drug Evaluation and Research, Silver Spring, MD. https://www.fda.gov/regulatory-information/search-fda-guidance-documents/statistical-approaches-establishing-bioequivalence

43. U.S. Food and Drug Administration. 2018. Bioanalytical method validation guidance for industry. Center for Drug Evaluation and Research, Rockville, MD. https://www.fda.gov/regulatory-information/search-fda-guidance-documents/bioanalytical-method-validation-guidance-industry

44. Houston JB. 1994. Utility of in vitro drug metabolism data in predicting in vivo metabolic clearance. Biochem Pharmacol 47:1469–1479. 10.1016/0006-2952(94)90520-7

45. Obach RS. 1999. Prediction of human clearance of twenty-nine drugs from hepatic microsomal intrinsic clearance data: an examination of in vitro half-life approach and nonspecific binding to microsomes. Drug Metab Dispos 27:1350–1359. 10.1016/s0090-9556(24)14938-0

46. Fluckiger-Isler S, Baumeister M, Braun K, Gervais V, Hasler-Nguyen N, Reimann R, Van Gompel J, Wunderlich HG, Engelhardt G. 2004. Assessment of the performance of the Ames II assay: a collaborative study with 19 coded compounds. Mutat Res 558:181–197. 10.1016/j.mrgentox.2003.12.001

47. OECD. 2020. Test no. 471: bacterial reverse mutation test. OECD guidelines for the testing of chemicals, section 4. OECD Publishing, Paris. 10.1787/9789264071247-en

48. OECD. 2016. Test no. 487: in vitro mammalian cell micronucleus test. OECD guidelines for the testing of chemicals, section 4. OECD Publishing, Paris. 10.1787/9789264264861-en

