## Supplementary Material for "MDL-001: an Oral, Direct-Acting Broad-Spectrum Antiviral for Influenza-Like Illness (ILI) and Chronic Hepatitis In Vivo"

### Supplementary Tables

***Supplementary Table S1. In vitro antiviral activity of MDL-001 by virus family, virus, and assay system.*** *Bold rows appear in Figure 1.*

| | Strain | Cell Line | $EC_{50}$ (µM) | $CC_{50}$ (µM) | $SI_{50}$ | Assay Type | Serum/protein condition: | Free Fraction $EC_{50}$ (nM) | Free Fraction $EC_{90}$ (nM) |
|---|---|---|---|---|---|---|---|---|---|
| ***Orthomyxoviridae*** | | | | | | | | | |
| **Influenza A** | **H1N1/California/04/09 (WT)** | **MDCK** | **0.165** | **15.1** | **91.2** | **Plaque (PFU)** | **0.3% BSA** | **41.3** | **47.3** |
| | **H3N2/Victoria/361/2011** | **MDCK** | **0.498** | **15.1** | **30.2** | **Plaque (PFU)** | **0.3% BSA** | **125** | **194** |
| **Influenza B** | **Victoria/Florida** | **MDCK** | **0.554** | **15.1** | **27.1** | **Plaque (PFU)** | **0.3% BSA** | **139** | **256** |
| | **Yamagata/Brisbane** | **MDCK** | **0.625** | **15.1** | **24.1** | **Plaque (PFU)** | **0.3% BSA** | **156** | **297** |
| ***Pneumoviridae*** | | | | | | | | | |
| **RSV** | **A2** | **A549** | **1.59** | **7.2** | **4.5** | **VYR (luciferase reporter)** | **10% FBS** | **79.4** | **127** |
| ***Coronaviridae*** | | | | | | | | | |
| **SARS-CoV-2** | **USA-WA1/2020** | **HeLa-ACE2** | **0.75** | **>50** | **>66.7** | **IF** | **2% FBS** | **188** | **345** |
| | **Alpha (B.1.1.7)** | **HeLa-ACE2** | **0.83** | **>50** | **>60.2** | **IF** | **2% FBS** | **208** | **505** |
| | **Beta (B.1.351)** | **HeLa-ACE2** | **1.05** | **>50** | **>47.6** | **IF** | **2% FBS** | **263** | **595** |
| | **Delta (B.1.617.2)** | **HeLa-ACE2** | **0.74** | **>50** | **>67.6** | **IF** | **2% FBS** | **185** | **423** |
| | **Omicron (B.1.1.529)** | **HeLa-ACE2** | **0.75** | **>50** | **>66.7** | **IF** | **2% FBS** | **188** | **345** |
| | Mouse Adapted-WA1 | HeLa-ACE2 | 0.712 | 7.652 | 10.7 | IF | 2% FBS | 178 | 300 |
| **α-coronavirus** | **229E** | **MRC-5** | **0.11** | **1.03** | **9.4** | **CPE** | **2% FBS** | **27.5** | **47.5** |
| | 229E | Huh7 | 0.42 | 8.4 | 20.0 | CPE | 2% FBS | 105 | NC |
| **β-coronavirus** | **OC43** | **A549-ACE2** | **0.49** | **>15** | **>30.6** | **CPE** | **2% FBS** | **123** | **330** |
| ***Flaviviridae*** | | | | | | | | | |
| **Hepatitis C** | **Genotype 1b (Con1)** | **Huh-luc/neo-ET** | **3.28** | **>10** | **>3.0** | **Replicon** | **10% FBS** | **164** | **460** |
| | **Genotype 2a (JFH-1)** | **Huh7.5.1** | **1.54** | **>10** | **>6.5** | **Replicon** | **10% FBS** | **77.0** | **300** |
| ***Hepadnaviridae*** | | | | | | | | | |
| **Hepatitis B** | **HepAD38** | **AD38** | **1.04** | **4.5** | **4.3** | **qPCR** | **10% FBS** | **51.8** | **87.5** |
| ***Kolmioviridae*** | | | | | | | | | |
| **Hepatitis D** | **HDV-1 (genotype I) HBV Co-infection** | **hNTCP-Huh7** | **2.13** | **>5** | **>2.3** | **Antigen ELISA** | **10% FBS** | **106** | **152** |

CPE: Cytopathic Effect, VYR: Viral Yield Reduction, IF: Immunofluorescence, NC: not calculated

***Supplementary Table S2. Lung log10 viral reduction from vehicle (Influenza Study).***

| Lung Burden (Day 5 p.i.) | Vehicle | Oseltamivir (TAM) (clinical eq. dose) | MDL-001 |
|---|---|---|---|
| $Log_{10}$ Reduction (mean) | — | 2.7 | 2.6 |
| 95% CI ($Log_{10}$ Reduction) | — | (2.6, 2.9) **** | (2.3, 2.8) **** |
| RNA copies/mg (geometric mean) | $3.2 \times 10^6$ | $5.9 \times 10^3$ | $8.9 \times 10^3$ |
| 95% CI (RNA copies/mg) | ($2.4 \times 10^6$, $4.4 \times 10^6$) | ($5.5 \times 10^3$, $6.3 \times 10^3$) | ($4.8 \times 10^3$, $1.6 \times 10^4$) |
| | Superiority Comparison | Non-Superior ($P = 0.14$) | |
| | Equivalency Comparison Difference 90% CI (MDL-001 - TAM) | Equivalent (−0.3 to 0.0 $log_{10}$) | |

*Geometric means and 95% confidence intervals are shown. (n=3/ group). Significance versus vehicle was tested by two-way ANOVA (****P < 0.0001). The MDL-001 versus clinical equivalent mouse dose of oseltamivir active-comparator superiority comparison was performed on animal-level absolute lung viral RNA reductions by two-sided unpaired Welch's unequal-variance t-test (P = 0.14). Equivalence was assessed by two one-sided tests (TOST) against a pre-specified margin of ±0.5 log10. See Methods.*

***Supplementary Table S3. Lung $log_{10}$ viral reduction from vehicle (SARS-CoV-2 Study).***

| **Lung Burden** | **Vehicle** | **MDL-001** | **Nirmatrelvir**<br>***Owen et al.* (16)**<br>**D4; $CCID_{50}$; n = 12** | **Molnupiravir**<br>***Jeong et al.* (17)**<br>**D5; $TCID_{50}$; n = 5** |
|---|---|---|---|---|
| **$Log_{10}$ Reduction (mean)** | — | 2.9<br>3 dp.i. | 1.4<br>4 d.p.i, $CCID_{50}$ | 1.0<br>5d.p.i, $TCID_{50}$ |
| **95% CI ($Log_{10}$ Reduction)** | — | (1.5, 4.3)<br>*** | SD = 0.648, n = 12 | SD = 0.508, n = 5 |
| **$TCID_{50}$/mL (geometric mean)** | $3.4 \times 10^7$ | $4.1 \times 10^4$ | **Comparison:** | |
| **95% CI ($TCID_{50}$/mL)** | $(4.2 \times 10^6, 2.7 \times 10^8)$ | $(2.4 \times 10^3, 7.0 \times 10^5)$ | **MDL-001 Superior**<br>($P = 0.023$) | **MDL-001 Superior**<br>($P = 0.008$) |

*Geometric means and 95% confidence intervals are shown for the MDL-001 study (n=9/group). Significance versus vehicle was tested by two-way ANOVA (***$P < 0.001$). Nirmatrelvir and molnupiravir values are literature comparator statistics extracted from Owen et al. and Jeong et al. Nirmatrelvir was measured at Day 4 p.i. by $CCID_{50}$, molnupiravir was measured at Day 5 p.i. by infectious-titer assay, and MDL-001 was measured at Day 3 p.i. by $TCID_{50}$. SD values in the literature columns are the variance inputs used for the Welch comparisons. Literature-comparator P values for superiority testing were calculated by two-sided unpaired Welch's unequal-variance t-tests on $Log_{10}$ reduction summary statistics as described in Methods.*

***Supplementary Table S4. Plasma log10 viral reduction from vehicle for each day (HCV Study).***

| Day | Sofosbuvir (clinical eq. dose) | MDL-001 | | |
|---|---|---|---|---|
| | $Log_{10}$ reduction | $Log_{10}$ reduction | | |
| **day 3** | -0.0 ± 0.1 | -0.1 ± 0.1 | | |
| **day 6** | 1.0 ± 0.1 | 0.8 ± 0.1 | | |
| **day 9** | 1.9 ± 0.1 | 1.7 ± 0.1 | | |
| **day 12** | 2.4 ± 0.2 | 2.0 ± 0.1 | | |
| **day 15** | 2.7 ± 0.2 | 2.3 ± 0.1 | | |
| **day 18** | 2.7 ± 0.1 | 2.6 ± 0.1 | **Superiority Comparison** | **Equivalence Comparison Difference 90% CI (MDL-001 - SOF)** |
| **day 21** | 2.8 ± 0.1 | 2.9 ± 0.1 | Non-Superior (*P* = 0.79) | Equivalent (0.0 to +0.2 $log_{10}$) |
| **day 24** | 3.0 ± 0.1 | 3.0 ± 0.1 | Non-Superior (*P* = 0.61) | Equivalent (−0.1 to +0.1 $log_{10}$) |
| **day 28** | 3.3 ± 0.2 | 3.3 ± 0.1 | Non-Superior (*P* = 0.75) | Equivalent (−0.2 to +0.1 $log_{10}$) |

*$Log_{10}$ reductions are shown as mean ± 95% CI. n = 6 per group. The MDL-001 versus sofosbuvir superiority comparisons were performed on animal-level plasma-viremia absolute reductions by two-sided unpaired Welch's unequal-variance t-tests (Day 21, P = 0.79; Day 24, P = 0.61; Day 28, P = 0.75). Equivalence was assessed by two one-sided tests (TOST) against a pre-specified margin of ±0.5 log10. See Methods.*

***Supplementary Table S5. Plasma HBsAg log10 reduction from vehicle for each day (HBV Study).***

| Day | Oral TAF | Oral MDL-001 | |
|---|---|---|---|
| **Day 1** | 0.0 ± 0.0 | 0.0 ± 0.0 | |
| **Day 2** | 0.1 ± 0.1 | 0.1 ± 0.0 | |
| **Day 3** | 0.2 ± 0.1 | 0.1 ± 0.0 | |
| **Day 4** | 0.3 ± 0.0 | 0.2 ± 0.0 | |
| **Day 5** | 0.3 ± 0.1 | 0.3 ± 0.0 | |
| **Day 6** | 0.4 ± 0.1 | 0.4 ± 0.0 | |
| **Day 7** | 0.5 ± 0.0 | 0.5 ± 0.0 | **Superiority Comparison** |
| **Day 8** | 0.6 ± 0.0 | 0.7 ± 0.0 | MDL-001 Superior (*P* = 0.0042) |
| **Day 9** | 0.7 ± 0.0 | 0.8 ± 0.0 | MDL-001 Superior (*P* = 0.0005) |
| **Day 10** | 0.8 ± 0.0 | 0.9 ± 0.0 | MDL-001 Superior (*P* = 0.0003) |
| **Day 13** | 1.0 ± 0.1 | 1.0 ± 0.0 | MDL-001 Superior (*P* = 0.0037) |
| **Day 16** | 1.1 ± 0.1 | 1.2 ± 0.0 | MDL-001 Superior (*P* = 0.0005) |
| **Day 19** | 1.2 ± 0.1 | 1.5 ± 0.1 | MDL-001 Superior (*P* <0.0001) |
| **Day 21** | 1.4 ± 0.2 | 1.7 ± 0.1 | MDL-001 Superior (*P* = 0.0059) |
| **Day 24** | 1.7 ± 0.3 | 1.9 ± 0.2 | MDL-001 Non-superior (P = 0.0746) |
| **Day 27** | **1.9 ± 0.4** | **2.3 ± 0.3** | LLOQ reached in 3/6 of MDL-001 mice, 1/6 of TAF mice |
| **Day 28** | **2.4 ± 0.1** | **2.5 ± 0.0 (LLOQ)** | **LLOQ** reached in all MDL-001 mice, 5/6 of TAF mice |

*$Log_{10}$ reductions are shown as mean ± 95% CI. n = 6 per group. The MDL-001 versus tenofovir alafenamide superiority comparisons were performed on log10-transformed per-animal values by two-sided unpaired Welch's unequal-variance t-tests. The assay lower limit of quantification was 1 ng/mL. Values at or below the limit were censored at the limit before transformation, so reductions on days when an arm reached the limit are lower bounds. See Methods.*

***Supplementary Table S6. Oral pharmacokinetics of MDL-001 in mouse plasma, lung, and liver after single oral doses of 25, 50, or 75 mg/kg in suspension.*** *Tissues were sampled at 1, 2, 4, 8, 16, and 24 h. Units are ng/mL for plasma and ng/g for tissues. Parameters are* $t_{max}$, $C_{max}$, $C_{24h}$, $AUC_{0\text{–}24h}$, *and* $t_{1/2}$. *NC indicates not calculated.*

| Oral PK of MDL-001 in Mouse Plasma | | | | |
|---|---|---|---|---|
| **Parameter** | **Units** | **Dose (mg/kg)** | | |
| | | 25 | 50 | 75 |
| $T_{max}$ | h | 2 | 1 | 1 |
| $C_{max}$ | ng/g | 122 | 155 | 280 |
| $C_{24h}$ | ng/g | 1.6 | 7.2 | 20.1 |
| $AUC_{0\text{-}24h}$ | ng-h/g | 670 | 937 | 2227 |
| $T_{1/2}$ | h | < 8 | NC | 8 |

| Oral PK of MDL-001 in Mouse Lung | | | | |
|---|---|---|---|---|
| **Parameter** | **Units** | **Dose (mg/kg)** | | |
| | | 25 | 50 | 75 |
| $T_{max}$ | h | 2 | 4 | 4 |
| $C_{max}$ | ng/g | 4722 | 8136 | 13348 |
| $C_{24h}$ | ng/g | 133 | 420 | 2310 |
| $AUC_{0\text{-}24h}$ | ng-h/g | 25729 | 67778 | 168258 |
| $T_{1/2}$ | h | 9 | NC | 8 |

| Oral PK of MDL-001 in Mouse Liver | | | | |
|---|---|---|---|---|
| **Parameter** | **Units** | **Dose (mg/kg)** | | |
| | | **25** | **50** | **75** |
| $T_{max}$ | h | 2 | 1 | 4 |
| $C_{max}$ | ng/g | 8636 | 16060 | 19870 |
| $C_{24h}$ | ng/g | 184 | 472 | 2414 |
| $AUC_{0\text{-}24h}$ | ng-h/g | 43746 | 85461 | 186764 |
| $T_{1/2}$ | h | 8 | 9 | 9 |

| **Lung Partition** | | **Dose (mg/kg)** | | |
|---|---|---|---|---|
| | | **25** | **50** | **75** |
| $C_{max}$-derived | $K_p$ | 39 | 52 | 48 |
| $AUC_{0\text{-}24h}$-derived | $K_p$ | 38 | 72 | 76 |

| **Liver Partition** | | **Dose (mg/kg)** | | |
|---|---|---|---|---|
| | | **25** | **50** | **75** |
| $C_{max}$-derived | $K_p$ | 71 | 104 | 71 |
| $AUC_{0\text{-}24h}$-derived | $K_p$ | 65 | 91 | 84 |

***Supplementary Table S7. Rat plasma pharmacokinetics of Oral MDL-001 formulation.*** *Non-compartmental pharmacokinetic parameters in male Sprague-Dawley rats following a single oral dose at 400 mg/kg (n = 3). Plasma was sampled at 0.5, 1, 2, 4, 6, 10, and 24 h post-dose.*

| ***Rat Plasma PK of MDL-001 (Optimized Oral Solution Formulation)*** | | |
|---|---|---|
| ***Parameter*** | ***Units*** | ***Dose (mg/kg)*** |
| | | ***400*** |
| ***Actual delivered dose*** | ***mg/kg*** | *543 ± 12.1* |
| ***$t_{max}$ (range)*** | ***h*** | *4 (2–10)* |
| ***$C_{max}$*** | ***ng/mL*** | *641 ± 18.5* |
| ***$C_{24h}$*** | ***ng/mL*** | *72.8 ± 67.9* |
| ***$t_{1/2}$*** | ***h*** | *NC* |
| ***$AUC_{last}$*** | ***h·ng/mL*** | *7150 ± 1470* |
| ***$AUC_{\infty}$*** | ***h·ng/mL*** | *NC* |

***Supplementary Table S8. Cross-Species In Vitro Intrinsic Clearance of MDL-001***

| **Species** | **$t_{½}$ (min)** | **$CL_{int,u}$ (µL/min/million)** | **In Vivo CL (mL/min/kg)** | **Hepatic Extraction Ratio (%)** | **Clearance Classification** |
|---|---|---|---|---|---|
| Mouse | 17.8 | 458 | 27.3 | 23 | Low |
| Rat | 18.5 | 440 | 22.2 | 33 | Intermediate |
| Dog | 16.9 | 484 | 7.97 | 26 | Low |
| Monkey | 22.4 | 364 | 2.13 | 5 | Low |
| Human | 53.2 | 153 | 1.68 | 8 | Low |

***Supplementary Table S9. Metabolite profile for MDL-001 across species in cryopreserved hepatocytes.*** *Human metabolites greater than 1.5% fraction are bold.*

| Metabolite | Identification | Human (%) | Mouse (%) | Rat (%) | Dog (%) | Monkey (%) | Buffer (%) |
|---|---|---|---|---|---|---|---|
| **MDL-001** | **Parent compound** | **33.7** | **6.6** | **31.9** | **8.4** | **7.5** | **95.5** |
| **M1** | **Oxidation in piperidine (N-oxide)** | **2.5** | **0.2** | **12.2** | **3.3** | **1.0** | **4.4** |
| M7 | Oxidative dealkylation + ox + dehydrogenation | 1.3 | 0.8 | 2.0 | 2.2 | 2.9 | – |
| **M10** | **Glucuronide conjugation (phenol)** | **45.0** | **31.1** | **8.0** | **12.5** | **72.1** | **–** |
| **M11** | **Glucuronide conjugation (phenol)** | **5.0** | **45.4** | **16.8** | **48.7** | **6.7** | **–** |
| M16 | Ox in piperidine N-oxide + glucuronide (phenol) | 0.9 | 0.6 | 1.4 | 0.8 | 1.4 | – |
| **M30** | **Sulfo-conjugation** | **7.9** | **0.1** | **3.4** | **3.1** | **0.9** | **–** |
| M31 | Ox in piperidine N-oxide + sulfo | 0.9 | – | 4.8 | 5.4 | 0.2 | – |

***Supplementary Table S10. Calculation of hERG safety margin from in vitro $IC_{50}$ and projected unbound human plasma $C_{max}$ at the minimally efficacious dose.***

| Parameter | Value |
|---|---|
| **hERG $IC_{50}$ (Predictor fluorescence-polarization assay; no plasma protein in incubation)** | 13 µM (13,000 nM) |
| **Rat plasma $C_{max}$ at a minimally efficacious dose** | 230 ng/mL |
| **Human plasma free fraction ($F_u$)** | 0.005 |
| **Projected unbound human plasma $C_{max}$ (= $C_{max} \times F_u$; converted to nM using MW 456.58 g/mol)** | 1.15 ng/mL = 2.5 nM |
| **hERG $IC_{50}$/ unbound $C_{max}$ safety margin** | >5,200-fold |

***Supplementary Table S11. Summary of duration, animal, and route of administration for the tolerability studies of MDL-001 in mice and rats (totaling 376 animals across seventeen experiments).***

| Study Duration | N (animals) | Route |
|---|---|---|
| 3 days | 6 mice | Oral |
| 3–4 days | 22 mice | Oral |
| 3 days | 36 mice | Oral |
| 10 days | 30 mice | Oral |
| 10 days | 18 mice | Oral |
| 28 days | 18 mice | Oral |
| 10–16 days | 24 mice | Oral |
| 28 days | 18 mice | Oral |
| 4–6 days | 18 mice | Oral |
| 4–7 days | 6 mice | Oral |
| 7 days | 3 mice | Oral |
| 1 dose | 90 mice | Oral |
| 5 days | 48 mice | Oral |
| 1 dose | 12 rats | Oral |
| 1 dose | 9 rats | Oral |
| 1 dose | 6 rats | IV |
| 1 dose | 12 rats | Oral |
